# Multi-omics Insight into Cardiac Myofibril Remodeling in Post-Prandial Burmese Pythons

**DOI:** 10.64898/2026.02.02.703272

**Authors:** Thomas G. Martin, Lorena Suarez-Artiles, Kathleen C. Woulfe, Elise G. Melhedegaard, Yuxiao Tan, Dakota R. Hunt, Bruce E. Kirkpatrick, Lia Nguyen, Joseph Lee, Isabella Laskey, Kristi S. Anseth, Julien Ochala, Michael Gotthardt, Philipp Mertins, Leslie A. Leinwand

## Abstract

Burmese pythons exhibit rapid cardiac remodeling in response to a dramatic increase in metabolic rate during digestion. Here, we performed single-myofibril mechanics measurements and myosin heavy chain metabolic assays to evaluate the impact of feeding on the cardiomyocyte sarcomere – the fundamental molecular unit of muscle contraction – using two experimental paradigms: normal feeding (one meal per month) and frequent feeding (eight meals per month). Myofibril tension and rate of relaxation increased during digestion in both paradigms, while frequent feeding was further associated with slower myofibril activation kinetics and faster myosin heavy chain ATP turnover. To identify molecular changes at the sarcomere and gain potential mechanistic insight, we performed multi-omics analyses. RNA sequencing identified increased expression of some sarcomere genes during digestion; however, proteomics analysis suggested a delay in sarcomere protein synthesis at the peak of remodeling, as expression of many sarcomere proteins decreased. Analysis of post-translational modifications (ubiquitinomics, phospho-proteomics, acetylomics) identified hundreds of significantly regulated sites on sarcomere proteins during digestion, including many on the tension-regulating titin and myosin heavy chain proteins. Our results detail the molecular underpinnings of cardiac remodeling in digesting Burmese pythons and suggest that nature’s solution for rapidly increasing cardiac contractility is a post-translational sarcomere tuning program.

## INTRODUCTION

Burmese pythons (*Python molurus bivittatus*) utilize an ambush-hunting predation strategy that is associated with consuming large prey across months-long intervals of fasting^1^. Consequently, they exhibit a >10-fold increase in metabolic rate during digestion, which coincides with increased size and activity of most organs, including the heart^2,3^. Cardiac adaptations in digesting Burmese pythons include modest, but rapid cardiac hypertrophy development (10-40% increased mass within 3 days), increased heart rate (∼2.5-fold), and increased ventricular stroke volume (2-fold)^2^ stemming from enhanced cardiomyocyte contractility^4^. These factors contribute to a ∼5-fold increase in cardiac output at the peak of digestion^2^, which is comparable to the change in the human heart going from rest to exercise^5^. The factorial increase in these parameters and rapid timescale of cardiac remodeling in Burmese pythons positions them as a useful model for deciphering the molecular regulation of cardiac plasticity^2,3^. Previous studies of Burmese pythons have identified important roles for hypertrophy-promoting circulating factors^6,7^, lipid regulation^8^, cardiomyocyte cell cycle re-entry^9^, and systemic oxygen supply-demand mismatch^10^ during adaptive cardiac remodeling. Nevertheless, the molecular mechanisms of cardiac functional changes in digesting Burmese pythons remain incompletely understood.

The smallest molecular unit of striated muscle contraction is the sarcomere^11^. A paracrystalline structure composed of hundreds of proteins, the sarcomere mediates calcium (Ca^2+^)-dependent shortening of muscle via tightly-regulated interactions between actin thin filaments and myosin thick filaments^11^. In mammals, sarcomere function can be tuned through changes in sarcomere protein isoform expression and/or post-translational modifications (PTMs), which contribute both to increased myofilament function with exercise^12,13^ and impaired function in heart failure^14,15^. Our recent study of Ball pythons (*Python regius*), a smaller snake species separated from Burmese pythons by ∼20 million years of evolution^16^, identified increased active (Ca^2+^-dependent) and decreased passive (Ca^2+^-independent) tension in cardiac myofibrils one day after feeding^17^. These results suggest that myofibril properties are modified to support enhanced cardiac function during digestion in pythons, but the enabling molecular changes at the sarcomere level were not investigated. Previous work in Burmese pythons identified increased mRNA expression of myosin heavy chain 6 (*MYH6*) post-prandially^18^, suggesting myosin isoform switching may contribute to myofibril functional changes, as the predominant myosin in the python heart is MYH15^19^. Another study identified increased expression of cardiomyocyte Ca^2+^-handling proteins that coincided with increased contractility and faster relaxation *ex vivo*^4^. Whether sarcomere modifications contribute to these post-prandial changes in contractile function remains unknown. We hypothesized that feeding engages coordinated, PTM-driven sarcomere adaptations that rapidly adjust myofibril mechanics to meet the transient metabolic demands of digestion.

In this study, we investigated the effects of feeding on single-myofibril mechanics, sarcomere gene and protein expression, and sarcomere protein PTMs in the Burmese python heart. We studied the responses to normal (one meal per month) and frequent (eight meals per month) feeding intervals, the latter of which induces sustained elevation of metabolism due to constant digestion^8,9^. Myofibril tension increased during digestion in both groups and this effect was more pronounced after frequent feeding. Digestion was also associated with increased rate of myofibril relaxation. Frequent feeding uniquely reduced myofibril activation rate and accelerated ATP turnover by myosin heavy chain.

Analysis of sarcomeric mRNAs by bulk RNA sequencing revealed increased expression of some genes after feeding, yet proteomics analysis identified a net decrease in relative sarcomere protein abundance, which coincided with decreased ventricular tissue density – consistent with an apparent dilution of sarcomere content during rapid tissue remodeling rather than a simple transcription-to-protein gain. Enrichment and quantification of PTMs (ubiquitinated lysine, phosphorylated serine/threonine/tyrosine, and acetylated lysine) by mass spectrometry identified hundreds of sarcomere modifications that were significantly altered during digestion. Dozens of modifications were identified on the tension-regulating titin and myosin heavy chain proteins, which may contribute to the observed changes in myofibril function with feeding. Altogether, our results highlight the dramatic sarcomere remodeling induced by feeding in Burmese pythons and suggest that enhanced cardiac contractility during digestion is accompanied by increased myofibril performance and extensive PTM remodeling of key mechano-active proteins.

## RESULTS

### Digestion is associated with increased myofibril active tension and altered activation and relaxation kinetics

Adult Burmese pythons were randomly assigned to receive 25% bodyweight rat meals in two different feeding intervals: normal feeding (one meal per month) and frequent feeding (FF, eight meals per month) (**Fig. 1A**). The frequently fed group was included to examine the effects of sustained metabolic demand and was inspired by the thriving, invasive population of pythons in the Florida Everglades where prey is abundant^18^. After two months of feeding in these intervals, pythons were humanely euthanized and cardiac ventricular tissue was collected. The timepoint of 3-days post-feeding (3DPF and FF-3DPF) was chosen for study as this had previously been identified as the peak of metabolic activity and organ remodeling post-prandially^2,3,6^. Fasted pythons (28DPF and FF-28DPF) were included as controls for both feeding groups.

**Fig. 1.**
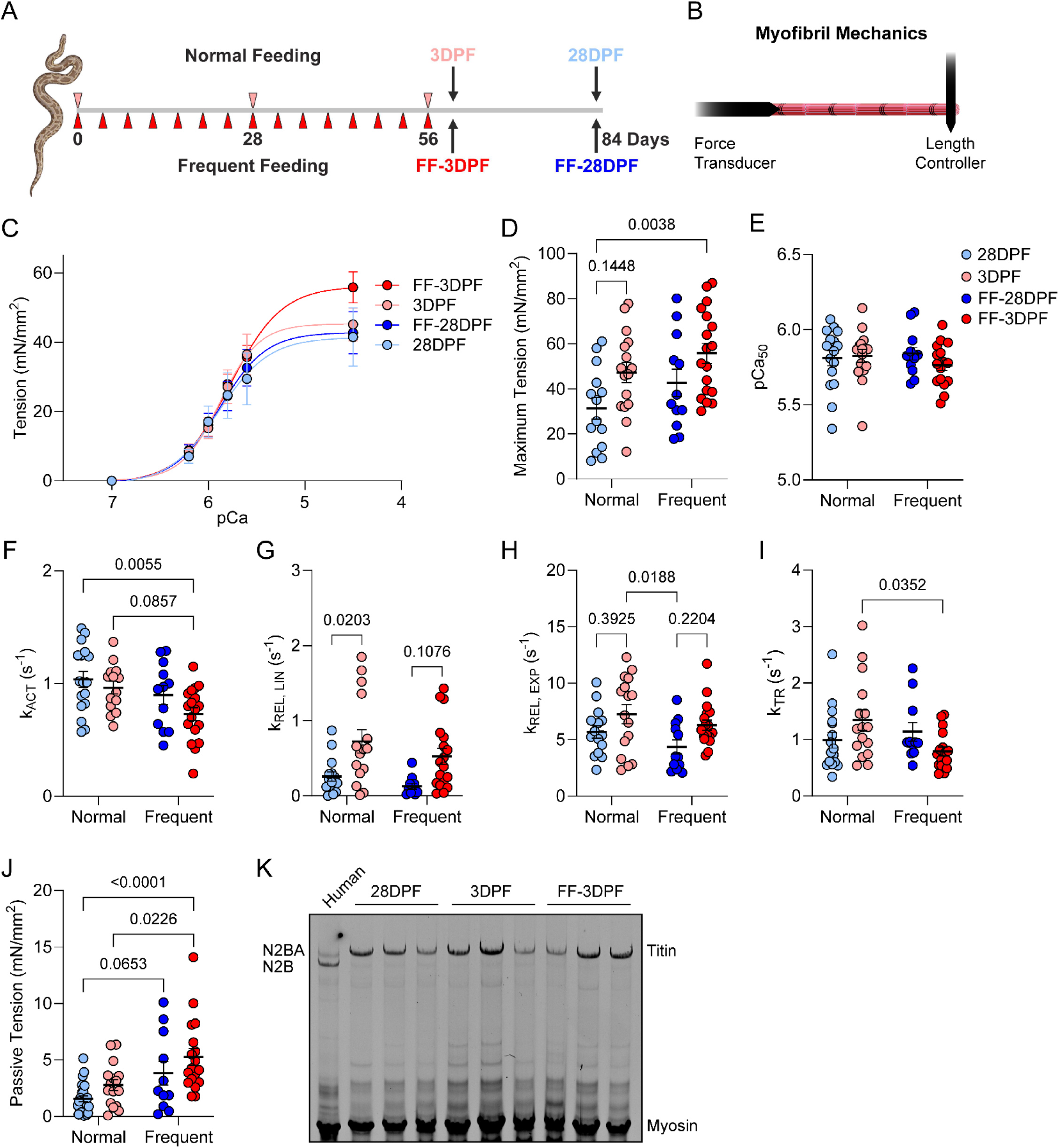
Digestion is associated with altered myofibril tension and kinetics. **A.** Experimental paradigm; made with BioRender. **B.** Schematic of myofibril functional assay setup. **C.** Myofibril tension-pCa curve; n = 18 28DPF, 17 3DPF, 12 FF-28DPF, and 18 FF-3DPF. **D.** Maximum tension normalized to cross-sectional area; n = 14 28DPF, 16 3DPF, 12 FF-28DPF, and 18 FF-3DPF; interaction: p = 0.782, feeding frequency: p = 0.049, time after feeding: p = 0.005. **E.** Myofibril pCa at 50% maximum tension (pCa_50_); n = 16 28DPF, 16 3DPF, 12 FF-28DPF, 16 FF-3DPF. **F.** Rate of myofibril activation (k_ACT_); n = 17 28DPF, 14 3DPF, 12 FF-28DPF, 18 FF-3DPF; interaction: p = 0.491, feeding frequency: p = 0.008, time after feeding: p = 0.077. **G.** Rate of the linear phase of relaxation (k_REL,LIN_); n = 15 28DPF, 15 3DPF, 10 FF-28DPF, 18 FF-3DPF; interaction: p = 0.763, feeding frequency: p = 0.148, time after feeding: p = 0.0003. **H.** Rate of the exponential phase of relaxation (k_REL,EXP_); n = 15 28DPF, 16 3DPF, 12 FF-28DPF, 18 FF-3DPF; interaction: p = 0.789, feeding frequency: p = 0.077, time after feeding: p = 0.008. **I.** Rate of tension redevelopment (k_TR_); n = 16 28DPF, 16 3DPF, 11 FF-28DPF, 17 FF-3DPF; interaction: p = 0.019, feeding frequency: p = 0.172, time after feeding: p = 0.983. **J.** Passive tension at sarcomere length 2.2 µm normalized to cross-sectional area; n = 26 28DPF, 16 3DPF, 11 FF-28DPF, 18 FF-3DPF; interaction: p = 0.854, feeding frequency: p = 0.0002, time after feeding: p = 0.031. **K.** Titin protein isoform gel for a human LV sample and python ventricular samples. For D-J, outliers determined by the ROUT outlier test were removed. Statistical analysis was performed by two-way ANOVA; if significance was detected, Sidak’s post-hoc test for multiple independent pairwise comparisons was employed. All data are presented as the mean ± SEM.

To determine the effects of digestion and feeding frequency on myofibril properties, single myofibrils were isolated, attached to a force transducer and high-speed length controller (**Fig. 1B**), and submitted to various functional tests. Myofibril tension and Ca^2+^ sensitivity were analyzed by force-pCa measurements (**Fig. 1C**). There was no interaction between feeding frequency and time after feeding for maximal tension by two-way ANOVA; however, tension was significantly affected by both feeding frequency (Frequent vs. Normal) and time after feeding (3DPF vs. 28DPF), and the increase in tension was more pronounced during digestion after frequent feeding (**Fig. 1D**). There was no difference in Ca^2+^ sensitivity between groups (**Fig. 1E**). Myofibril rate of activation (*k*_ACT_) was significantly affected by feeding frequency but not time after feeding, where frequent feeding led to decreased *k*_ACT_ (**Fig. 1F**). Meanwhile, the rates of the linear (*k*_REL,LIN_) and exponential (*k*_REL,EXP_) phases of relaxation both increased as functions of time after feeding but not feeding frequency (**Fig. 1G-H**), indicating a general functional response to feeding. For the rate of tension redevelopment (*k*_TR_), which represents the sum of myosin cross-bridge attachment and detachment rates^20^, there was a significant interaction between feeding frequency and time after feeding with *k*_TR_ decreased in FF-3DPF versus 3DPF (**Fig. 1I**).

### Feeding increases myofibril passive tension independent of titin isoform changes

At fixed sarcomere lengths, myofibril force-pCa and activation/relaxation kinetics describe Ca^2+^-dependent properties and are predominantly driven by thin filament Ca^2+^-responsiveness and myosin cross-bridge cycling. However, Ca^2+^-independent properties also play an important role in regulating sarcomere function. Passive tension describes the elastic force generated by the myofibril at a given sarcomere length in the absence of Ca^2+^ and is primarily due to the spring-like sarcomere protein titin (interchangeable with TTN hereafter)^21,22^. Passive tension, measured in a pCa 9 solution at a sarcomere length of 2.2 µm, increased as a function of time after feeding and feeding frequency, but the effect was more pronounced during digestion after a frequent feeding interval (**Fig. 1J**). Notably, while other myofibril parameters were restored by 28 days after frequent feeding, passive tension remained trending higher in FF-28DPF versus 28DPF (p = 0.065, **Fig. 1J**). Analysis of stiffness and elastic properties at the ventricular tissue level by rheology revealed that stiffness (increased) and elasticity (decreased) were significantly affected by increased feeding frequency, irrespective of time after feeding (**Fig. S1A-B**). These data suggest that myofibril stiffness changes induced by frequent feeding translate to the organ level, although the potential contribution of ECM remodeling at the tissue-scale cannot be ruled out. In humans and other mammals, increased myofibril passive tension can result from expression of shorter titin isoforms^23^ or changes in stiffness-modifying titin post-translational modifications^24^. To determine whether feeding affected titin isoform expression, we performed large format agarose gel electrophoresis to separate titin isoforms (**Fig. 1K**). Compared to the human LV, which displayed the expected N2BA (3.3 MDa) and N2B (3.0 MDa) titin isoforms, Burmese python ventricular samples had only one dominant titin isoform that migrated more slowly than human N2BA (∼3.4 MDa) (**Fig. 1K**). Therefore, we concluded that changes in myofibril passive tension with feeding were not due to altered titin isoform expression. Together with the above, these results highlight the general responses to feeding (increased tension and *k*_REL_) and unique impact of increased feeding frequency (decreased *k*_ACT_ and *k*_TR_) on myofibril properties.

### Frequent feeding decreases myosin heavy chain ATP turnover time without affecting DRX:SRX ratio

Myofilament force generation is directly proportional to the number of actin-myosin cross-bridges formed^25^. Myosin can exist in at least three distinct states: actin-bound, disordered relaxed (DRX), and super-relaxed (SRX)^26^. The relative proportion of myosins in the DRX (fast ATP turnover) versus SRX (slow ATP turnover) states contributes significantly to cardiomyocyte contractility and energetics^27^. To determine whether digestion impacted myosin DRX:SRX ratio, we pulsed skinned ventricular strips from 28DPF, 3DPF, and FF-3DPF pythons with fluorescently labeled ATP (Mant-ATP) and then chased with dark ATP to assess turnover kinetics (**Fig. 2A**). The FF-28DPF group was not analyzed as no myofibril functional differences were identified between this group and 28DPF. Fluorescence in this assay decays according to a two-phase exponential model from which DRX and SRX states can be deconvoluted (**Fig. 2A**)^28^. In fasted pythons, ∼40% of myosins were in the DRX state, similar to the proportion in hibernating mammals^29^, and this was not altered during digestion after a normal feeding interval (**Fig. 2B**). Digestion after a frequent feeding interval was associated with a slight, but not statistically significant, increase in DRX myosin (**Fig. 2B-C**). While the proportion of myosin in these states was unchanged, the time constant of ATP turnover in the DRX state (T1) significantly decreased in FF-3DPF versus 3DPF (**Fig. 2D**). The SRX myosin time constant (T2) also trended toward a decrease in FF-3DPF (p = 0.102, **Fig. 2E**). Together, the changes in DRX and SRX ATP turnover times led to a trending increase (p = 0.089) in relative ATP consumption by myosin in FF-3DPF versus 3DPF (**Fig. 2F**). These data indicate that myosin biochemical states are not affected during digestion after a normal feeding interval, while frequent feeding modestly decreases DRX myosin ATP turnover time.

**Fig. 2.**
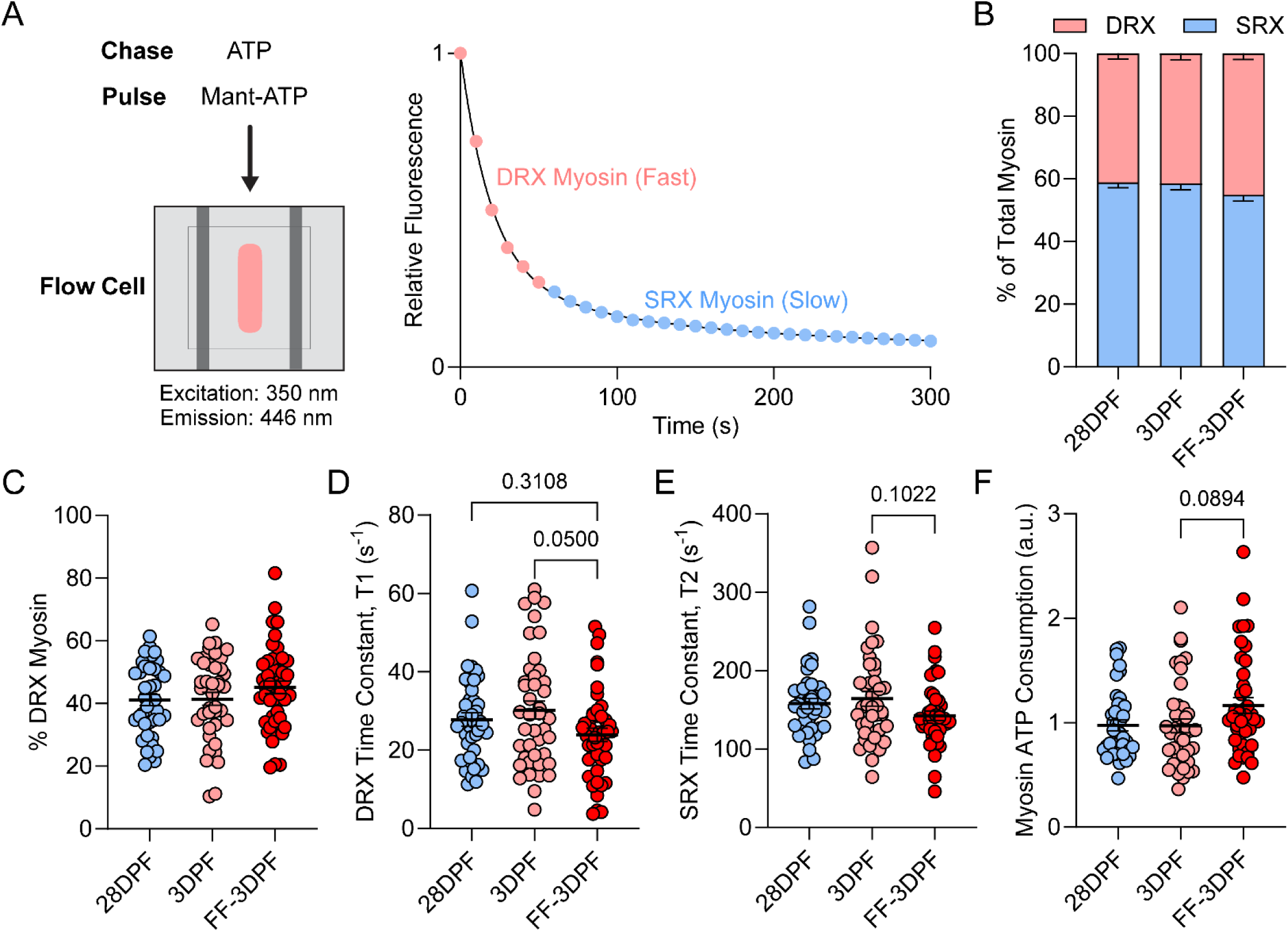
Frequent feeding induces faster ATP turnover by myosin heavy chain. **A.** Schematic of the experimental setup for the Mant-ATP pulse-chase assay and a representative graph displaying the two-phase exponential decay of fluorescence attributed to DRX and SRX myosin. **B.** Proportion of myosin in the DRX and SRX states in skinned cardiac ventricular tissue strips. **C.** Proportion of myosin in the DRX state. **D.** Time constant of ATP turnover for myosin in the DRX state (T1). **E.** Time constant of ATP turnover for myosin in the SRX state (T2). **F.** Relative myosin ATP consumption represented as a fold-change vs. the 28DPF group. For all, n = 41 28DPF, 42 3DPF, 45 FF-3DPF. Statistical analysis was performed by one-way ANOVA with Tukey’s post-hoc test for multiple independent pairwise comparisons. Data are presented as the mean ± SEM.

### Increased sarcomere gene expression with feeding is insufficient to maintain sarcomere protein density at the peak of remodeling

To gain insight into molecular regulation of myofibril function during digestion, we began by measuring sarcomere gene and protein expression. First, we performed bulk mRNA sequencing to assess transcriptome remodeling in response to feeding. Principal component dimensionality reduction analysis (PCA) revealed that the transcriptional profiles of the four groups clustered based on time after feeding and were not markedly affected by feeding frequency (**Fig. 3A**). Digestion after each type of feeding was associated with ∼3,000 differentially expressed genes (**Fig. 3B-C**). Gene Ontology (GO) Biological Process analysis identified protein translation, RNA processing, and autophagy among the downregulated processes in both feeding paradigms, while protein folding and cell metabolism increased after a normal feeding interval (**Fig. 3D-F**). Frequent feeding was uniquely linked to increased expression of mitosis and cell cycle genes (**Fig. 3G, Fig. S2**), matching previous observations^9^. Focusing on genes encoding sarcomere proteins, 17 were significantly differentially expressed at 3DPF (13 up, 4 down) (**Fig. 3H**) and *MYH6* was among genes with increased expression, matching previous observations^18^. Expression of 12 sarcomere genes changed after frequent feeding (7 up, 5 down), although here *MYH6* decreased (**Fig. 3I**). *MYH6, MYH7*, *MYL9, FHL3*, and *PDLIM7* gene expression was significantly increased in 3DPF versus FF-3DPF hearts (**Fig. 3J**).

**Fig. 3.**
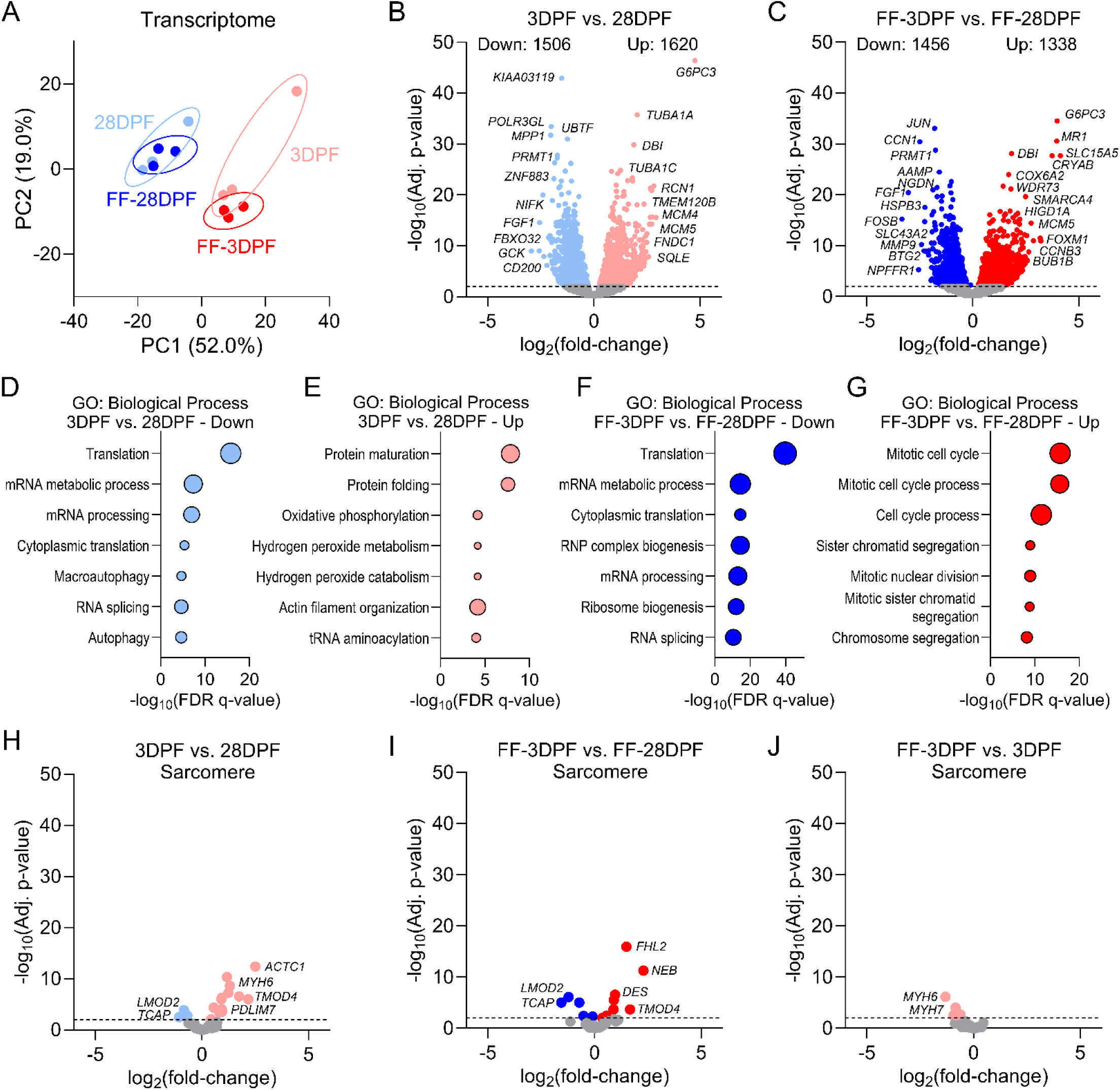
Feeding induces differential expression of sarcomere genes. **A.** PCA plot for bulk RNA sequencing data from post-fed Burmese python cardiac ventricular samples; n = 3/group. **B-C.** Volcano plots depicting fold-change, statistical significance, and number of significantly differentially expressed genes in the 3DPF vs. 28DPF (B) and FF-3DPF vs. FF-28DPF (C) comparisons. **D-E**. GO: Biological Process enrichment for the downregulated (D) and upregulated (E) genes in the 3DPF vs. 28DPF comparison. **F-G.** GO: Biological Process enrichment for the downregulated (F) and upregulated (G) genes in the FF-3DPF vs. FF-28DPF comparison. **H-J.** Volcano plots showing only the sarcomere genes from our mRNA sequencing analysis and depicting fold-change and statistical significance for sarcomere genes in the 3DPF vs. 28DPF (H), FF-3DPF vs. FF-28DPF (I), and FF3DPF vs. 3DPF (J) comparisons.

To determine if transcriptome-level differences translated to the proteome, we performed label-free data independent acquisition (DIA) quantitative proteomics. PCA of the proteomics data showed no overlap between the four groups (**Fig. 4A**). Over 800 and 2,000 proteins were differentially expressed during digestion after normal and frequent feeding intervals, respectively (**Fig. 4B-C**). GO Biological Process analysis of the down- and up-regulated proteins identified decreased cellular respiration and increased protein synthesis with both feeding intervals (**Fig. 4D-G, Fig. S3**). Focusing on the sarcomere, we found 12 differentially expressed proteins, most of which (11) decreased during digestion (**Fig. 4H**). Digestion after frequent feeding was associated with more robust sarcomere protein expression changes, although here too the majority (31/35) were found to decrease (**Fig. 4I**). Recent work in the post-prandial Ball python heart identified increased cytoplasmic area by electron microscopy, suggesting reduced myofibril density during digestion^17^. To determine whether ventricular tissue density decreased post-prandially, we loaded tissue samples with a fluorescent molecule (calcein) and then bleached the signal with a high-powered laser and measured fluorescence recovery over time to monitor diffusion rate through the tissue. We found a faster diffusion rate in the 3DPF and FF-3DPF groups indicating decreased tissue density during digestion (**Fig. 4J**). These data support that, while force per sarcomere increases during digestion (myofibril mechanics), tissue sarcomere density decreases.

**Fig. 4.**
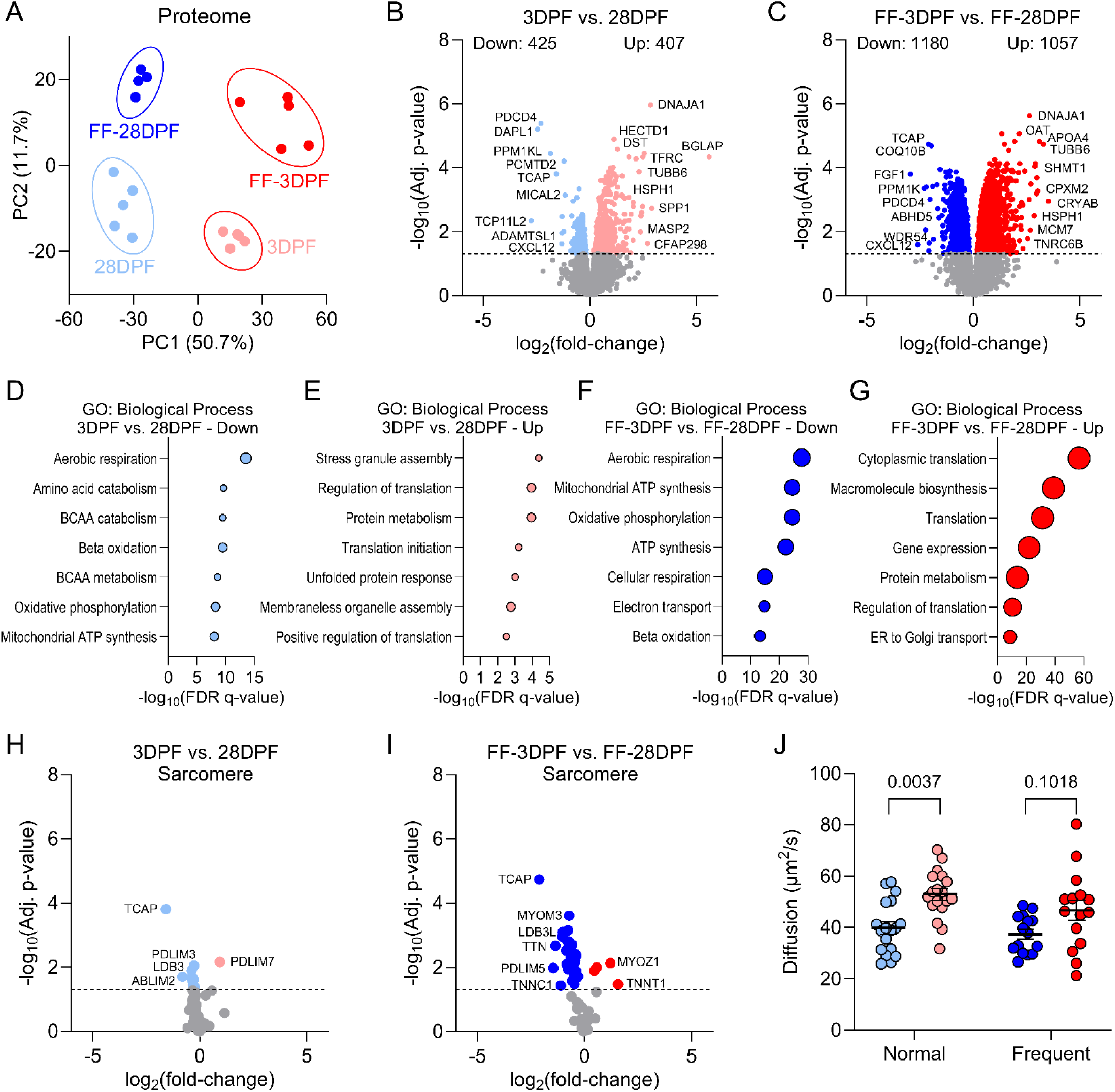
Relative sarcomere protein abundance and cardiac tissue density decrease post-feeding. **A.** PCA plot for quantitative proteomics data from post-fed Burmese python cardiac ventricular samples; n = 4-5/group. **B-C.** Volcano plots depicting fold-change, statistical significance, and number of significantly differentially expressed proteins in the 3DPF vs. 28DPF (B) and FF-3DPF vs. FF-28DPF (C) comparisons. **D-E**. GO: Biological Process enrichment for the downregulated (D) and upregulated (E) proteins in the 3DPF vs. 28DPF comparison. **F-G.** GO: Biological Process enrichment for the downregulated (F) and upregulated (G) proteins in the FF-3DPF vs. FF-28DPF comparison. **H-I.** Volcano plots depicting fold-change and statistical significance showing only the sarcomere proteins from the 3DPF vs. 28DPF (H) and FF-3DPF vs. FF-28DPF (I) comparisons. **J.** Calcein fluorescence recovery after photobleaching (FRAP), denoting the diffusion rate of the molecule through python ventricular tissue (faster recovery/increased diffusion rate = decreased tissue density); n = 15-18 samples from 3 biological replicates per group; two-way ANOVA with Tukey’s post-hoc test: interaction: p = 0.492, feeding frequency: p = 0.114, time after feeding: p = 0.0001.

### Feeding-associated ubiquitinome changes largely track with protein abundance

We reasoned that the disconnect between sarcomere transcript and protein abundance could be due to two logical possibilities: 1. New sarcomere protein synthesis was unable to match the rate of tissue growth, despite increased gene expression (i.e., there was a delay in protein translation); 2. Sarcomere proteins were being actively removed through protein degradation pathways. We hypothesized that, given the rapid timescale of cardiac remodeling in pythons, the former was likely true. However, to examine the second possibility, we performed ubiquitin-enrichment proteomics (ubiquitinomics). Ubiquitin is a post-translational modification that marks proteins for degradation by the ubiquitin-proteasome and autophagy-lysosome systems^11^. Our analysis revealed that ubiquitination generally increased during digestion at 3DPF and protein synthesis pathways were among the top enriched processes (**Fig. 5A-B**). Comparison of ubiquitination site fold-changes with total protein abundance showed a strong positive correlation (**Fig. 5C**). Digestion after a frequent feeding interval coincided with more potent remodeling of the ubiquitinome, including decreased ubiquitination at the sarcomere and increased ubiquitination of protein synthesis factors (**Fig. 5D-F**). In most cases, the decreased ubiquitination of sarcomere proteins tracked with their total protein abundance; however, sites with increased ubiquitination were found on some proteins with reduced total protein abundance (TTN, MYH15, and FLNC) (**Fig. 5G**). GO Biological Process analysis of proteins with decreased abundance and at least one increased ubiquitination site identified enrichment of plasma membrane and sarcomere components (**Fig. 5H**). No significant ubiquitination sites were identified between FF-3DPF and 3DPF (**Fig. 5I**). These findings suggest that, in most cases, the disconnect between sarcomere mRNA and protein at the peak of cardiac remodeling is due to a lag between transcription and accumulation of stable sarcomere protein mass during rapid growth (i.e., synthesis does not immediately keep pace with remodeling).

**Fig. 5.**
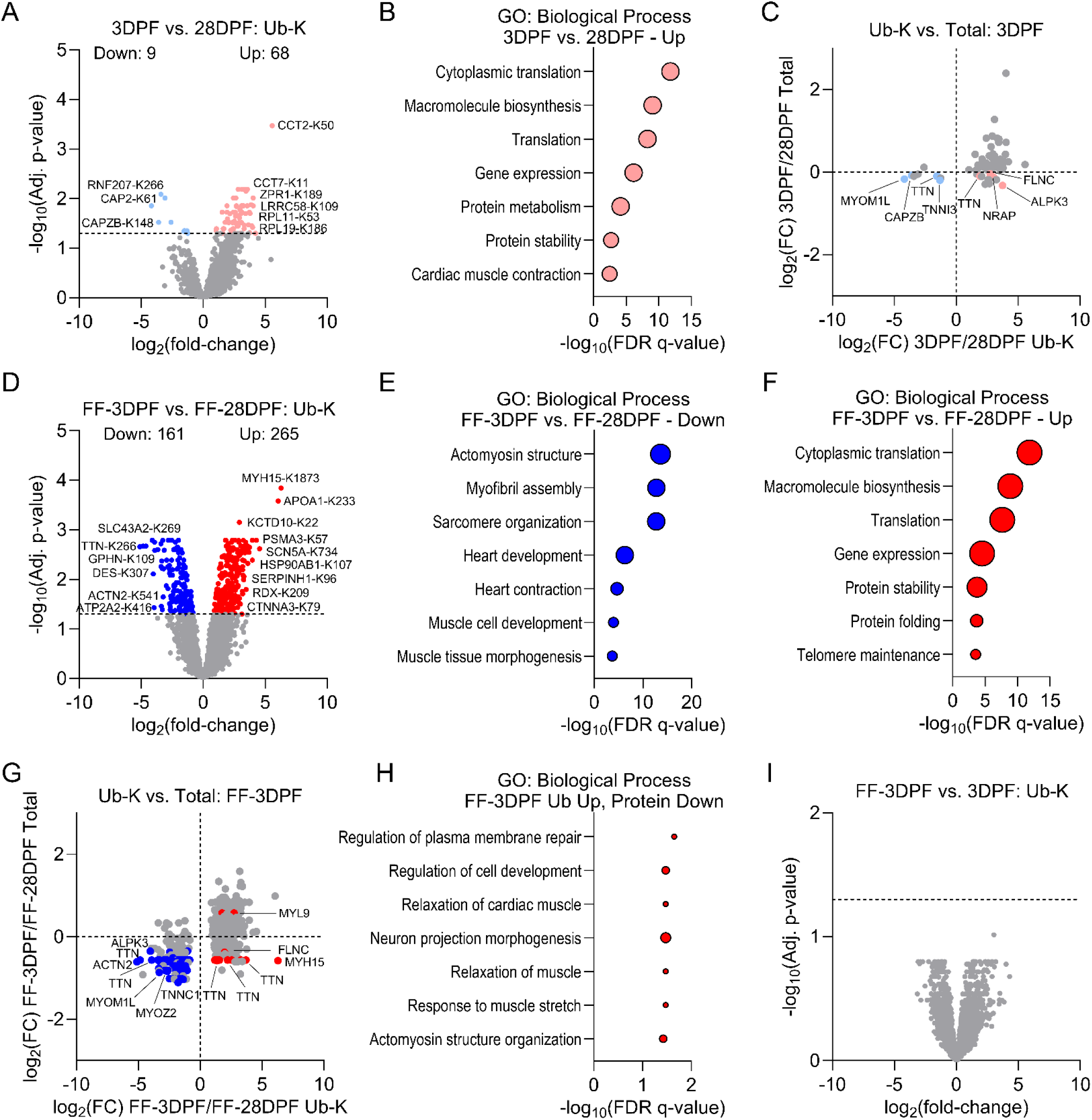
Cardiac ubiquitinome changes with feeding track with relative protein abundance. **A.** Volcano plot depicting fold-change, statistical significance, and number of significantly differentially ubiquitinated sites at 3DPF vs. 28DPF. **B.** GO: Biological Process enrichment for the upregulated ubiquitination sites. **C.** Total protein fold-change vs. ubiquitination site fold-change for the significantly differentially abundant ubiquitination sites at 3DPF vs. 28DPF; sarcomere proteins are annotated. **D.** Volcano plot depicting fold-change, statistical significance, and number of significantly differentially ubiquitinated sites at FF-3DPF vs. FF-28DPF. **E-F.** GO: Biological Process enrichment for the downregulated (E) and upregulated (F) ubiquitination sites. **G.** Total protein fold-change vs. ubiquitination site fold-change for the significantly differentially abundant ubiquitination sites at FF-3DPF vs. FF-28DPF; sarcomere proteins are annotated. **H.** GO: Biological Process enrichment of the proteins with increased ubiquitination and decreased total abundance. **I.** Volcano plot depicting fold-change and significance for ubiquitination sites at FF-3DPF vs. 3DPF.

### The sarcomere phospho-proteome is dramatically remodeled post-prandially

Decreased sarcomere protein density with digestion can contribute to altered tissue-level contractility; however, it cannot explain functional changes at the single myofibril level where sarcomere density is fixed. Therefore, we next analyzed post-translational modifications that have previously been described to modulate sarcomere function: phosphorylation^24,30–32^ and acetylation^24,33–35^. Enrichment of phospho-serine/threonine/tyrosine peptides and analysis by mass spectrometry (phospho-proteomics) revealed that, based on PCA, the digestive responses were largely similar regardless of feeding interval (**Fig. 6A**). Digestion in both paradigms was associated with potent remodeling of the phospho-proteome with 3,638 and 6,911 significantly regulated sites identified in 3DPF and FF-3DPF, respectively (**Fig. 6B-C**). Over 2,500 phospho-sites were uniquely regulated during digestion after frequent feeding (**Fig. 6D**) and 161 remained significant between FF-28DPF and 28DPF (**Fig. S4**). To deconvolute changes in phosphorylation from total protein abundance changes, we analyzed phospho-site fold-change versus total protein fold-change and selected only those sites with >1 or <-1 log_2_ fold-change at the phospho-site level versus total protein for enrichment analyses (**Fig. 6E-F**). In both feeding intervals, GO Biological Process analysis identified decreased phosphorylation of nuclear pore proteins post-feeding, while myofibril proteins were enriched among both the down- and up-regulated phospho-sites (**Fig. 6G-J**). PTM signature enrichment analysis identified shared kinase activation signatures associated with digestion in both paradigms (CK2A2, MAPK12, and ERK2) and others that were specific to the frequent feeding interval (DYRK4, JNK1, MARK2) (**Fig. 6K**). That unique kinase activation patterns contribute to distinct sarcomere remodeling was supported by the identification of over 350 significantly differentially regulated sarcomere phospho-sites between FF-3DPF and 3DPF (**Fig. S5A**). Among sites with increased phosphorylation at FF-3DPF were multiple MYH15 and TTN residues (**Fig. S5B-D**), which may contribute to the observed feeding frequency-dependent differences in myofibril function.

**Fig. 6.**
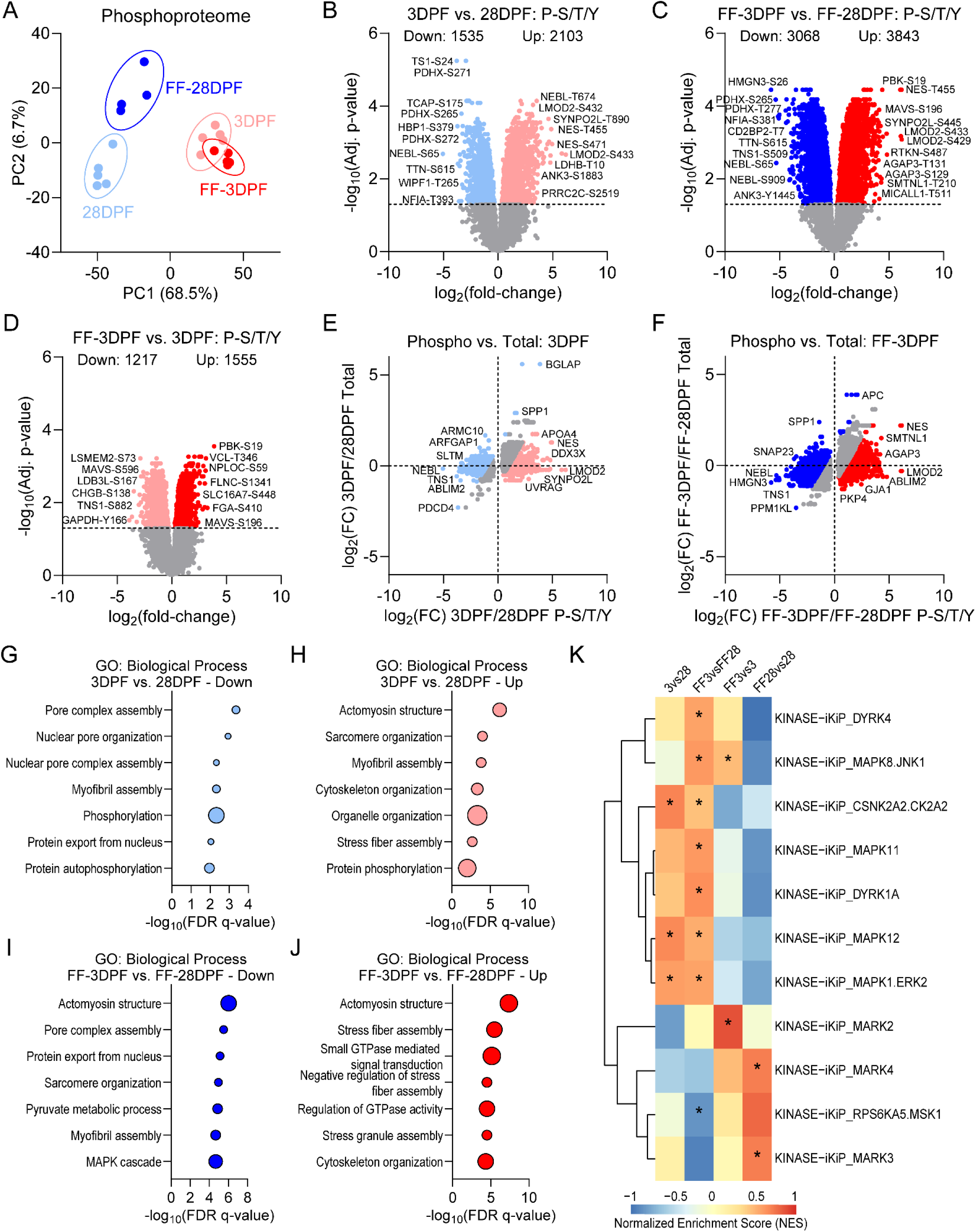
Feeding dramatically remodels the cardiac ventricular phospho-proteome. **A.** PCA plot for phospho-proteomics data from post-fed Burmese python cardiac ventricular samples; n = 4-5/group. **B-D.** Volcano plots depicting fold-change, statistical significance, and number of significantly differentially expressed phospho-sites in the 3DPF vs. 28DPF (B), FF-3DPF vs. FF-28DPF (C), and FF-3DPF vs. 3DPF comparisons. **E-F.** Total protein fold-change vs. phospho-site fold-change for the significantly differentially abundant phospho-sites at 3DPF vs. 28DPF (E) and FF-3DPF vs. FF-28DPF (F). **G-H**. GO: Biological Process enrichment for the phospho-sites that decreased by log_2_ fold-change ≥ 1.0 (G) or increased by log_2_ fold-change ≥ 1.0 (H) vs. total protein abundance in the 3DPF vs. 28DPF comparison. **I-J.** GO: Biological Process enrichment for the phospho-sites that decreased by log_2_ fold-change ≥ 1.0 (G) or increased by log_2_ fold-change ≥ 1.0 (H) vs. total protein abundance in the FF-3DPF vs. FF-28DPF comparison. **K.** Heat map depicting normalized enrichment scores (NES, columns) of signatures (rows) for significant in vitro kinase-to-phosphosite (iKiP) database hits in the different post-feeding group comparisons by ptmSEA; asterisks in heat map denote statistical significance.

### Feeding induces increased protein acetylation, including multiple sites on titin and myosin

To determine the effects of digestion and feeding frequency on the acetylome, we enriched and analyzed acetylated lysine residues by mass spectrometry. This analysis identified 85 (75 up, 10 down) significant sites at 3DPF versus 28DPF (**Fig. 7A**). Proteins with increased acetylation included factors involved in protein acetylation and chromatin organization (**Fig. 7B**), suggesting increased chromatin accessibility during digestion, matching previous observations in Ball pythons^17^. Significantly increased acetylation sites were identified on the sarcomere proteins TTN, MYH15, and PDLIM5-like (**Fig. 7C**). Digestion after a frequent feeding interval was associated with even further remodeling of the acetylome (104 up, 25 down) (**Fig. 7D**). Here too, most proteins with increased acetylation were involved in chromatin remodeling and protein acetylation (**Fig. 7E**); however, three acetylated lysine residues in TTN and one in MYH15 were also identified that were similarly regulated in 3DPF. Most of the downregulated acetylation sites in FF-3DPF were on sarcomere proteins, likely reflecting their reduced total protein abundance in this group (**Fig. 7F**). Next, we assessed the impact of the different feeding intervals on ventricular acetylome remodeling to identify modifications uniquely regulated by frequent feeding (**Fig. 7G**). This analysis identified three sites on sarcomere proteins (MYH15 Ac-K1395, ACTN2 Ac-K792, and MYOZ2 Ac-K172) that were significantly affected by feeding frequency but not time after feeding (**Fig. 7H-J**). Acetylation at these sites remained elevated even with a subsequent 28-day fast (**Fig. 7H-J**), highlighting the long-term sarcomere post-translational remodeling induced by frequent feeding.

**Fig. 7.**
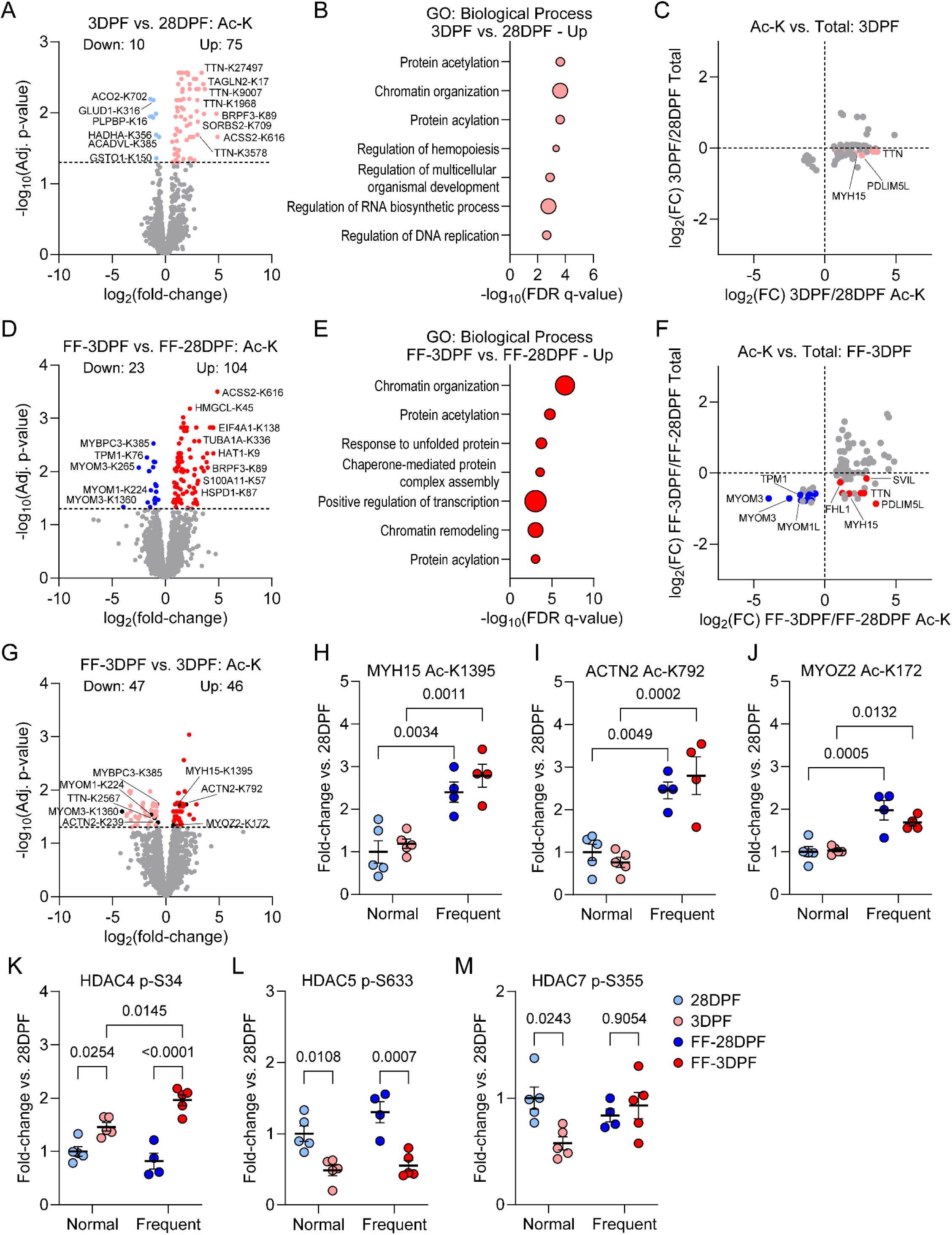
Digestion is associated with sarcomere acetylome changes. **A.** Volcano plot depicting fold-change, statistical significance, and number of significantly differentially acetylated sites at 3DPF vs. 28DPF. **B.** GO: Biological Process enrichment for the upregulated acetylation sites at 3DPF. **C.** Total protein fold-change vs. acetylated lysine site fold-change for the significantly differentially abundant acetylation sites at 3DPF vs. 28DPF; sarcomere proteins are annotated. **D.** Volcano plot depicting fold-change, statistical significance, and number of significantly differentially acetylated sites at FF-3DPF vs. FF-28DPF. **E.** GO: Biological Process enrichment for the upregulated acetylation sites at FF-3DPF. **F.** Total protein fold-change vs. acetylated lysine site fold-change for the significantly differentially abundant acetylation sites at FF-3DPF vs. FF-28DPF; sarcomere proteins are annotated. **G.** Volcano plot depicting fold-change, statistical significance, and number of significantly differentially acetylated sites at FF-3DPF vs. 3DPF. **H-J.** Examples differentially acetylated sites uniquely modified according to feeding frequency. **K-M.** Phosphorylation sites regulated after feeding on Histone deacetylase-4, -5, and -7. For H-M, the data were analyzed by two-way ANOVA with Tukey’s post-hoc test for multiple independent pairwise comparisons and are presented as the mean ± SEM.

Notably, protein acetylation after both feeding paradigms generally increased, suggesting global activation of acetyltransferases and/or inactivation of deacetylases during digestion. This observation agrees with prior work in Ball pythons where cardiac histone deacetylase activity was found to decrease post-prandially^17^. Interestingly, our phosphoproteomics data revealed significant changes in phosphorylation shared between feeding paradigms for HDAC4 and -5, but not HDAC7 (**Fig. 7K-M**). According to the PhosphoSite Plus database, S34 in HDAC4 and S633 in HDAC5 are conserved in humans (S265 and S661, respectively), and changes in their phosphorylation have been linked to HDAC subcellular localization^36,37^. Of note, S633 phosphorylation of HDAC5 is reduced after feeding, and PKD-dependent phosphorylation of this site regulates 14-3-3 binding, and subsequent cytoplasmic retention, of HDAC5 in neonatal rat ventricular myocytes^36^. It has been shown that cAMP induces hypo-phosphorylation of 14-3-3 binding sites in HDAC5, leading to nuclear accumulation and subsequent inhibition of MEF2 in cardiomyocytes^38^. Thus, we hypothesize that the post-prandial increase in acetylation of sarcomere proteins, including TTN and MYH15, could be a consequence of HDAC-MEF2 axis regulation in response to feeding-induced hypertrophy.

### Summary of modifications on the tension-regulating titin and myosin heavy chain proteins

Considering the entirety of sarcomere PTM changes during digestion, our analyses identified dozens of modifications on TTN that were shared between feeding groups (**Fig. 8A**) and over 100 more that were unique to frequent feeding (**Fig. 8B**). Notably, the Z-disk and M-band regions of TTN were hotspots for modifications, although PTMs were found throughout the molecule, including many in the I-band region, which generates most of TTN-based passive tension^39^. We identified six feeding-regulated modifications on MYH15 that were shared between feeding groups, two unique to 3DPF, and dozens more exclusively in FF-3DPF (**Fig. 8C-D**). Most of these modifications were found in the rod domain, which regulates sarcomere incorporation^40^; however, several were also identified in the N-terminal motor domain responsible for ATP-dependent cross-bridge cycling^11^. Some PTMs on TTN and MYH15 were found in domains that have previously been implicated in tension generation, including the PEVK domain, immunoglobulin (Ig)-like, and fibronectin-3 (Fn3)-like domains in the I-band region of TTN (**Fig. 8E**)^24,41^ and the converter domain of MYH15 (**Fig. 8F**)^42^. Our findings serve as a starting point for future mechanistic investigations into the specific myofibril functional consequences of these modifications.

**Fig. 8.**
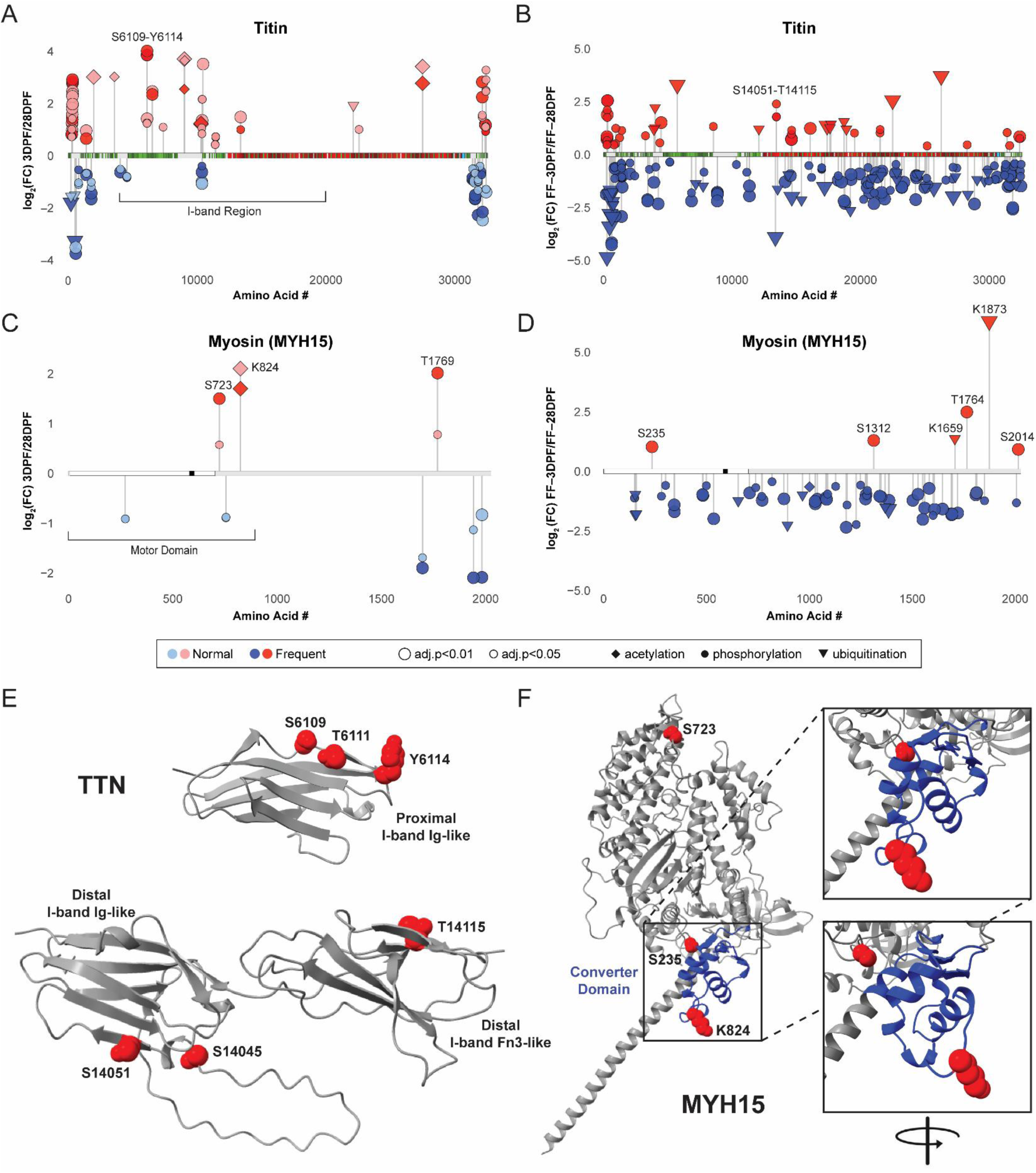
Feeding-induced post-translation modifications on titin and myosin heavy chain. **A.** Summary of increased and decreased PTMs on titin that were shared in both feeding groups. **B.** Summary of titin PTMs that were uniquely regulated in the frequent feeding group. **C.** Summary of increased and decreased PTMs on myosin heavy chain (MYH15) that were shared in both feeding groups. **D.** Summary of MYH15 PTMs that were uniquely regulated in the frequent feeding group. **E.** AlphaFold3 structural predictions and amino acid residue annotations for phosphorylation sites on titin I-band immunoglobulin (Ig)-like and fibronectin-3 (Fn3)-like domains that were particularly enriched after frequent feeding. ‘Proximal’ and ‘Distal’ denotations are in reference to the Z-disk. **F.** AlphaFold3 structural predictions and amino acid residue annotations for PTM sites in the MYH15 motor domain that were altered by feeding: S723 and K824 in 3DPF and FF-3DPF; S235 only in FF-3DPF.

## DISCUSSION

Digesting Burmese pythons are a model of extreme cardiometabolic plasticity^3^. Compared to mammalian models of adaptive cardiac remodeling, which develop hypertrophy and enhanced cardiac function over weeks^43^, the heart of digesting pythons reaches its peak structural and functional remodeling within 72 hours of feeding^2,3^. The rapid timescale and magnitude of remodeling position Burmese pythons as a valuable model for investigating the molecular regulation of adaptive cardiac plasticity^2^. Previous studies of the heart in post-prandial Burmese pythons identified changes in cardiac physiology^10,44–46^, mechanisms of cardiac hypertrophy and regression^6,7,10^, transcriptional remodeling^9,47–51^, and increased *ex vivo* cardiac contractility^4^. However, cardiac functional remodeling at the fundamental molecular scale of the sarcomere had not been assessed, leaving much to be discovered regarding their digestion-associated adaptations. In this study, we investigated the impact of feeding on sarcomere function and found that digestion was associated with increased myofibril tension and rate of relaxation, while frequent feeding additionally altered myofibril activation kinetics and myosin heavy chain ATP turnover. Furthermore, we comprehensively characterized the changes in sarcomere protein expression and PTMs that coincided with these functional differences via multi-omics analyses, lending potential molecular insight into mechanisms for tuning myofibril function. Our data support that nature’s solution for rapidly modulating cardiac contractility without inducing pathology is a post-translational sarcomere tuning program.

While sarcomere-level insights into cardiac remodeling in Burmese pythons were lacking, we had previously investigated myofibril function in digesting Ball pythons at 24 hours post-feeding after a normal feeding interval^17^. As observed in the present study, digestion in Ball pythons coincided with increased myofibril maximum Ca^2+^-dependent tension^17^. However, whereas we observed increased myofibril relaxation rate in Burmese pythons at 72 hours, this property was significantly reduced at 24 hours in Ball pythons^17^. Also opposite to the findings presented herein, Ball python myofibrils displayed decreased passive tension with feeding^17^. These differences in functional adaptations to digestion may reflect the divergence between these species with respect to meal size, metabolic rate regulation, and growth potential. Ball pythons are much more limited in terms of growth, reaching lengths of up to 1.8 meters compared to 6 meters for Burmese. In our experience, Ball pythons are also unlikely to eat prey larger than 25% of their bodyweight while Burmese pythons are capable of consuming prey equaling their own size^1,52^. The resulting different maximum metabolic requirements may explain species differences in cardiac adaptations with feeding. Future investigations into digestion-associated sarcomere PTMs in Ball pythons will be valuable to help explain the opposite myofibril relaxation kinetics and passive tension responses to those observed in Burmese pythons. Additionally, an assessment of myofibril properties across a range of post-feeding timepoints in both species will be informative.

In this study, we identified increased maximum myofibril tension independent of changes in Ca^2+^ sensitivity. Since Ca^2+^ sensitivity is primarily mediated by thin filament proteins^53^, this result implicates altered thick filament properties as the underlying mechanism. Myofibril active tension represents the ensemble of forces produced by hundreds of individual myosin molecules within sarcomeres. Therefore, mechanisms that increase cross-bridge number (e.g., increased myosin DRX state) and efficiency (e.g., optimal filament overlap and decreased lattice spacing), promote greater myosin step sizes (the distance a myosin head moves actin per ATP), or increase duty ratio (the time myosin heads spend bound to actin) represent possible explanations^54–60^. While we cannot rule out the contribution of changing sarcomere lengths to the increased *in vivo* cardiac contractility during digestion, our single myofibril studies were performed at fixed sarcomere lengths. Thus, differences in optimal filament overlap or lattice spacing (titin-dependent properties)^61^ are not expected to underlie the observed differences in active tension. Our analysis of DRX/SRX myosin states also suggests that myosin head availability is not a major contributor. Thus, changes in myosin intrinsic properties (e.g., intrinsic force and/or step size) represent a logical mechanism for the active tension increase. The converter domain of myosin heavy chain is crucial for transmitting force and regulating step size^62–64^. Mutations in the converter domain of MYH7 (the predominant cardiac myosin in humans) are also causative for hypertrophic cardiomyopathy (HCM)^63^, a disease associated with hypercontractility, and biopsies from patients with MYH7 converter domain mutations display increased maximum tension^65,66^. Our analysis of digestion-associated PTMs identified modifications in (Ac-K824, homologous to human MYH7 Q734) and structurally adjacent (p-S235, homologous to human MYH7 S148) to the Burmese python MYH15 converter domain. Intriguingly, Q734 lies in a region that is a hotspot for HCM-causing mutations in human MYH7^67,68^ and a missense mutation at S148 is listed as “likely pathogenic” for HCM in ClinVar^69^. MYH15 K824 acetylation increased during digestion irrespective of feeding interval, which may help explain the general increase in maximum tension observed as a function of time after feeding. Meanwhile, S235 phosphorylation was unique to frequent feeding, possibly contributing to the more pronounced active tension increase observed in this group. The potential to reversibly modulate myosin function through PTMs has obvious translational potential for human heart disease therapies. Thus, future studies investigating the functional effects of the myosin motor domain PTMs identified herein will be valuable.

The increased myofibril passive tension identified during digestion was particularly pronounced after a frequent feeding interval and coincided with changes in ventricular tissue stiffness and elasticity. In mammalian myofibrils at the same sarcomere length (2.2 µm), passive tension arises from sarcomere components (i.e., titin), while non-myofibril structures (e.g., microtubules) only make significant contributions at longer sarcomere lengths^70,71^. Based on this, we expect the increased passive tension during digestion in Burmese python myofibrils is due to changes in titin stiffness. Our study shows that python myofibrils contain only a single major titin isoform, unlike mammals which have two. Thus, changes in titin isoform expression cannot explain the phenotype. Titin PTMs are also known to contribute to passive tension^24^ and we identified hundreds of significantly altered PTMs on titin during digestion. The effects of most titin PTMs on tension are unknown. However, in mammals, phosphorylation at two regions has been linked to stiffness regulation: the N2B-unstructured region (phosphorylation = decreased tension) and the region immediately following the PEVK domain (phosphorylation = increased tension)^24^. Further, a recent study identified that acetylation of titin in the lysine-rich PEVK domain increases passive tension^72^. The Burmese python N2B domain is poorly conserved with human making inferences about PTMs in this region difficult; however, we did identify increased phosphorylation at multiple PEVK-adjacent sites (including T10373, S10375, T10870, T10876) and increased acetylation at three sites within the PEVK domain (K9007, K9039, K10301). In total, over 30 PTMs were found to increase in the I-band region of titin during digestion with roughly half of these being unique to the frequent feeding paradigm. A single PTM cannot confer the passive tension changes observed during digestion, but the ensemble effect of many modifications together in and near the PEVK may offer an explanation. Future single molecule atomic force microscopy studies investigating these and other titin modifications identified to ascertain their collective impact on tension will be valuable.

Taken together, our findings highlight the potent myofibril functional remodeling that occurs during digestion in the Burmese python heart and provide an atlas of feeding-regulated sarcomere PTMs. We also identified potential mechanisms responsible for these changes, including postprandial activation of kinase pathways (CK2A2, MAPK12, and ERK2) and suggested altered subcellular localization of deacetylases (HDAC4 and HDAC5). The combination of these datasets provides a launchpad for future mechanistic investigations that interrogate the individual and collective contributions of sarcomere protein PTMs to cardiac functional regulation and prioritizes specific, testable candidate sites on titin and MYH15. A key limitation is that our multi-omics analyses are associative; establishing causality will require targeted perturbation of prioritized PTM sites (e.g., site-directed mutagenesis or enzymatic modulation in recombinant/isolated systems) coupled to single-molecule and myofibril functional assays. Toward this end, we believe that examinations of myosin and titin properties will be most tractable, due to the already-developed assays for studying these proteins at a single molecule level^19,41^. Finally, we chose to focus our analysis of the multi-omics data on the sarcomere. However, there is a wealth of information regarding transcriptome and proteome remodeling present in these datasets that future researchers interested in other aspects of cardiac plasticity regulation may find valuable.

## METHODS

### Ethical Oversight

All animal experiments were performed with strict adherence to institutional and national guidelines for the care and use of laboratory animals. Procedures involving animals were conducted at the University of Colorado Boulder after approval by the Institutional Animal Care and Use Committee (IACUC). The University of Colorado Boulder Office of Animal Resources is an AAALAC International accredited facility.

### Burmese Python Husbandry and Experimentation

Captive-bred Burmese pythons (*Python molurus bivittatus*) were acquired as juveniles from Bob Clark Reptiles, Oklahoma City, Oklahoma. The pythons were singly housed at the University of Colorado Boulder in a room maintained at 30°C with 12-hour light-dark cycles. Pythons weighed ∼2 kg and were randomly assigned to treatment groups at the start of the study. For the normal feeding frequency groups, pythons were fed two 25% bodyweight rat meals at 28-day intervals. After the second feeding, pythons were euthanized by isoflurane overdose and rapid decapitation at 3- or 28-days post-feeding (DPF), and their cardiac ventricle tissue was collected, snap frozen in liquid nitrogen, and stored at -80°C until downstream processing and analysis. For the frequent feeding groups, pythons were fed a 25% bodyweight meal once every four days for eight weeks. After the final feeding, pythons were euthanized at 3- and 28-days post-frequent feeding (FF-3DPF, FF-28DPF). Changes in the mass of the heart and other organs in these cohorts in response to feeding were reported previously^9^.

### Myofibril Isolation

Myofibrils were isolated as previously described^17^. Frozen python ventricle tissues were skinned in Linke’s solution (132 mM NaCl, 5 mM KCl, 1 mM MgCl_2_, 10 mM Tris-base, 5 mM EGTA, 1 mM sodium azide, pH 7.0) with 1% Triton X-100 and protease inhibitors (10 μM leupeptin, 5 μM pepstatin, 200 μM phenyl-methylsuphonyl fluoride (PMSF), 10 μM E64, 500 μM NaN_3_, 2 mM dithioerythritol) for 1 hour at 4°C. Skinned myofibrils were washed in rigor solution (50 mM Tris, 100 mM KCl, 2 mM MgCl_2_, 1 mM EGTA, pH 7.0) then placed in bath solution (100mM Na_2_EGTA; 1M potassium propionate; 100 mM Na_2_SO_4_; 1M MOPS; 1M MgCl_2_; 6.7 mM ATP; and 1 mM creatine phosphate; pH 7.0) with protease inhibitors then homogenized (Tissue-Tearor).

### Myofibril Functional Measurements

Myofibril mechanical function was assessed on the day of isolation using the fast solution switching method as previously described^17,73^. Briefly, a droplet of myofibril suspension was added to a temperature-controlled chamber (15°C) and relaxing solution added. Small bundles of myofibrils (average diameter of 4.90 µm and average length of 73.9 µm) were mounted between two microtools (a calibrated cantilevered force probe (9.11 μm/μN) and a microtool attached to a motor that produces rapid length changes). Myofibril sarcomere length was set at ∼2.2 µm. Mounted myofibrils were activated and relaxed by rapidly translating the interface between two flowing streams of solutions (pCa 4.5 – maximum calcium concentration, and pCa 9.0 – relaxation solution). Data were collected and analyzed using customized LabView software.

### Large Format Agarose Gel Electrophoresis

The protocol for separation and visualization of titin protein isoforms was performed as previously described^74^. Briefly, 20 mg of frozen myocardium was pulverized using a 2 mL Dounce homogenizer submerged in liquid nitrogen. After tempering at -20°C for 10 minutes, the samples were solubilized in 1:40 (w/v) 8M urea, 2M thiourea buffer, 30% glycerol buffer and incubated at 60°C for 10 minutes. Samples were clarified by centrifugation, snap frozen, and stored at -80°C. A 1.5 mm thick 1% Seakem Gold agarose (Lonza) gel was assembled using the Hoefer SE6015 system. The gel was run on the day it was made. Protein samples were thawed at room temperature and 7.5 µL loaded per well. The gel apparatus was then moved to a container with cold running buffer (SDS-Tris-Glycine) in a 4°C cold room and run at 15 mA constant current for 3 hours and 15 minutes. The gel was removed from the glass plates and fixed in 50% methanol, 12.5% glacial acetic acid, 5% glycerol buffer overnight with gentle agitation and then stained with 0.05% (w/v) Coomassie brilliant blue for 3 hours. Destaining was performed with 25% methanol, 10% acetic acid for 5 hours. Imaging was performed using an Amersham Typhoon 5 imager (Cytiva) using the Coomassie in IR setting.

### Mant-ATP Chase Experiments

As previously published^28^, the relaxing solution contained 4 mM Mg-ATP, 1 mM free Mg^2+^, 10-6.0 µM free Ca^2+^ (pCa 9.0), 10 mM imidazole, 7 mM EGTA, 14.5 mM creatine phosphate. Ionic strength was adjusted to 180 mM and pH to 7.0. Additionally, the rigor buffer for Mant-ATP chase experiments contained 120 mM K acetate, 5 mM Mg acetate, 2.5 mM K_2_HPO4, 50 mM MOPS, 2 mM DTT with a pH of 6.8. On the day of the experiments, thin cardiac strips were isolated. Their ends were individually clamped to half-split copper meshes designed for electron microscopy (SPI G100 2010C-XA, width, 3 mm), which had been glued to glass slides (Academy, 26 x 76 mm, thickness 1.00-1.20 mm). Cover slips were attached to the top using double-sided tape to create flow chambers (Menzel-Gläser, 22 x 22 mm, thickness 0.13-0.16 mm)^28^. Subsequently, at 22°C (or 4°C), for each strip, the sarcomere length was measured using the brightfield mode of a Zeiss Axio Scope A1 microscope, a Zeiss AxioCam ICm 1 camera and a LD Plan-Neofluar 40x/0.60 objective. Preparations with a sarcomere length of 1.9-2.0 µm were further subjected to the protocol. Indeed, similar to previously published protocols^28^, each thin strip was first incubated for five minutes with a rigor buffer. A solution containing the rigor buffer with 250 μM Mant-ATP ((2’-(or-3’)-O-(N-Methylanthraniloyl) Adenosine 5’-Triphosphate) was then flushed and kept in the chamber for five minutes. At the end of this step, another solution made of the rigor buffer with 4 mM ATP was added with simultaneous acquisition of the Mant-ATP chase. For fluorescence acquisition, a Zeiss Axio Scope A1 microscope was used with a Plan-Apochromat 20x/0.8 objective and a Zeiss AxioCam ICm 1 camera. Frames were acquired every five seconds with a 20 ms exposure time using a DAPI filter set, for 5 min. Tests were run prior to starting the current study where images were collected for 15 min instead of 5 min – these tests did not reveal any significant difference in the parameters calculated. Three regions of each individual strip were sampled for fluorescence decay using the ROI manager in ImageJ as previously published^28^. The mean background fluorescence intensity was subtracted from the average of the fiber fluorescence intensity (for each image taken). Each time point was then normalized by the fluorescence intensity of the final Mant-ATP image before washout (T = 0). These data were then fit to an unconstrained double exponential decay using GraphPad Prism (version 10.4):

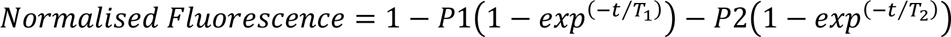

Where P1 is the amplitude of the initial rapid decay approximating the DRX state with T1 as the time constant for this decay. P2 is the slower second decay approximating the proportion of myosin heads in the SRX state with its associated time constant T2. In the present article, P1, P2, T1 and T2 are presented without any correction for the rapid wash-out of non-specific binding of Mant-ATP^28^. Solely based on P1, P2, T1, and T2, we estimated the theoretical ATP consumed by myosin^28^.

### Fluorescence Recovery after Photobleaching (FRAP)

FRAP was performed as previously described^75^. Briefly, tissue was equilibrated in a solution of calcein (2 µM in water) for at least 30 minutes before the sample was placed on a laser scanning confocal microscope (Nikon A1R). At least three regions of interest from separate imaging planes were selected in each tissue, with baseline fluorescence intensity (λ_ex_ = 488 nm) approximately 60% of the detection limit of the microscope. Following bleaching with 405 nm laser light at 100% power, fluorescence recovery was monitored for 90 seconds and fit to a closed formula in MATLAB to calculate the characteristic timescale of diffusion.

### Tissue Rheology

The modulus of ventricles was measured with oscillatory shear rheology (1 rad/s, 1% strain). A shear rheometer (DHR-3, TA Instruments) equipped with a sandblasted parallel plate geometry was used to measure the storage (G’) and loss (G’’) moduli and tan delta (G’’/G’) of the ventricle tissue at room temperature; tissue samples were compressed to an axial force of approximately 0.1 N before measurement. Young’s modulus was calculated by multiplying G’ by a value of 3 (i.e., assuming a Poisson ratio of 0.5, representing tissue incompressibility)^76^. Prior to measurement, frozen python ventricle tissues (n = 3) were thawed on ice and cut to size using an 8 mm cylindrical punch. Ventricle tissue was measured within 1 hour of defrosting. Tissue was trimmed to ensure a level height of ∼1 mm, and sandpaper was affixed to the Peltier plate to prevent tissue slippage.

### RNA Isolation

Cardiac ventricle tissue (∼50 mg) was added to 5 mL polypropylene tubes containing 1 mL of Trizol and homogenized using a mechanical homogenizer (Omni International). Samples were then incubated for five minutes at room temperature and transferred to 1.5 mL Eppendorf tubes. Chloroform added at 1:5 (vol/vol), the tubes shaken vigorously, and incubated at room temperature for 15 minutes prior to centrifugation at 12,000 RCF for 15 minutes at 4°C. The upper aqueous layer (∼500 µL) was collected into a clean tube, isopropanol added at 1:1 (vol/vol), and the samples briefly vortexed. To facilitate increased RNA precipitation, the tubes were incubated at -20°C for 30 minutes. The samples were then centrifuged at 12,000 RCF for 10 minutes to pellet the RNA. The supernatant was decanted, and the RNA pellet was washed twice with ice-cold 75% ethanol by resuspension and centrifugation. Following the last ethanol wash, the sample was air-dried for ∼10 minutes at room temperature and then the RNA was dissolved in 200 µL milli-Q water.

### RNA Sequencing

Poly-A RNA enrichment, cDNA library preparation, and short-read Illumina mRNA sequencing were performed by Novogene Corporation (Sacramento, CA) with the Illumina NovaSeq X system. Sample quality was assessed using FastQC version 0.11.51 and reads in FASTQ format were cleaned up using Trimmomatic version 0.36^77^ with the following parameters: ILLUMINACLIP:/opt/trimmomatic/0.36/adapters/TruSeq3-PE.fa:2:30:10 LEADING:3 TRAILING:3 SLIDINGWINDOW:4:15 MINLEN:36.Trimmed paired-end reads were mapped to the *Python molurus bivittatus* reference genome (NCBI RefSeq #GCF_000186305.1) using HISAT2 version 2.1.0^78^ with the --sensitive and --dta flags. Samtools version 1.8^79^ was used to convert SAM files to sorted and indexed BAM files. The Rsubread version 2.0.1^80^ featureCounts function was used to generate raw counts for each sample using the indexed BAM files and the reference annotation GTF file. Gene count normalization and differential gene expression analysis was conducted with DESeq2 version 1.36.0^81^. Pairwise comparisons between each group were conducted and the normal shrinkage estimator was used to produce shrunken log fold changes for visualization. Differentially expressed genes were defined as those with an adjusted p-value < 0.01. Over-representation analysis was performed on differentially expressed genes identified from RNA-Seq data. A gene annotation database for *Python bivittatus* was obtained through the AnnotationHub^82^ (v3.14.0) and AnnotationDbi^83^ (v1.68.0) packages in R. The complete set of genes tested for differential expression served as the background gene set. Functional enrichment of Gene Ontology (GO) biological processes was conducted separately for upregulated and downregulated gene sets using the clusterProfiler package (v4.14.4)^84^. Over-representation analysis was performed with the enrichGO function using the Benjamini–Hochberg method for multiple testing correction, and cutoffs of p < 0.05 and q < 0.1. Minimum and maximum gene set sizes were restricted to 10 and 500, respectively.

### Proteomics Sample Preparation

Frozen left ventricle tissue samples were cryo-pulverized using the Covaris CP02 CryoPrep Pulverizer with an impact level of 1. Pulverized samples were transferred to 2 mL Eppendorf tubes and resuspended in 500 µL of chilled 8M urea lysis buffer (8 M urea in water, 75 mM NaCl, 50 mM Tris pH 8.0, and 1 mM EDTA, supplemented with 2 µg/mL Aprotinin, 10 µg/mL Leupeptin, 1 mM PMSF, 10 mM NaF, 1:100 phosphatase-inhibitor cocktails 2 and 3, and 5 mM CAA). Samples were mixed by vortex for 10 seconds and incubated on ice for 15 minutes; this step was repeated twice. Tubes were centrifuged at maximum speed for 10 minutes to remove cell debris, and the supernatants were transferred to new 2 mL Eppendorf tubes. Protein concentration was estimated using BCA assay, and 1.5 mg protein aliquots were taken for in-solution digestion. Denatured proteins were reduced by adding a final concentration of 5 mM DTT and incubating for 1 hour at 37°C, and alkylated by incubation in the dark with 10 mM iodoacetamide (IAA) for 45 minutes at 25°C. Samples were diluted 1:4 with 50 mM Tris HCl pH 8.0 to decrease the Urea concentration below 2M. Proteins were pre-digested with LysC (1:50 enzyme to substrate ratio) for 2 hours at 25°C and digested overnight with Trypsin (1:50, 25°C). Peptides were acidified by adding formic acid (FA) to a final concentration of 1% and desalted using SepPak C18 columns (Waters) following the manufacturer’s instructions. For PTM analysis, 100 µg of peptides were used for IMAC phosphopeptide enrichment automated on an AssayMap Bravo System (Agilent) equipped with AssayMAP Fe(III)-NTA cartridges. 1 mg of peptides were used for serial ubiquitination and acetylation enrichment following the UbiFast ubiquitin profiling protocol^85^ using the PTMScan® HS Ubiquitin/SUMO Remnant Motif (K-ε-GG) Kit and the PTMScan® HS Acetyl-Lysine Motif (Ac-K) Kit (Cell Signaling Technology, 59322S and 46784S).

### Label-free Proteomics Analysis

Dried, desalted peptides were reconstituted in MS sample buffer containing 3% acetonitrile (ACN)/0.1% formic acid (FA) in water. For global proteome analysis 1 µg of peptides was injected into the system. One sixth or half of the enrichment output was used for phosphoproteome or ubiquitinome and acetylome analysis respectively. Samples were measured using a Vanquish Neo UHPLC system in nanoflow mode coupled with an Orbitrap Exploris 480 mass spectrometer (Thermo Fisher Scientific). Peptides were separated on a 20-cm reverse-phase column (inner diameter 75 μm, packed in-house with 1.9 μm C18 Reprosil beads) at a flow rate of 250 nL/min at 55° C. Mobile phase A was 0.1% (vol/vol) FA and 3% (vol/vol) ACN in water, while mobile phase B was 90% (vol/vol) ACN and 0.1% (vol/vol) FA in water. The gradient was increased from 2% to 90% B in 99 min, followed by a plateau at 90% B for 5 minutes. MS data were acquired in data-independent acquisition (DIA) mode. The spray voltage was set to 2.2 kV in positive ion mode. The expected LC peak was 10 seconds. Full scan MS spectra were acquired at a resolution of 120K in a scan range of 350-1650 m/z, 55% RF lens, 300% normalized automated gain control (AGC) target, 20 milliseconds maximum injection time (IT). DIA scans were done at a maximum injection time of 54 milliseconds at 30K resolution, 12 m/z isolation window, 26, 29, and 32 % stepped normalized collision energy (NCE), and 3000% normalized AGC target.

### Raw Proteomics Data Processing

Raw data were processed using Spectronaut 18 software (Biognosys) with the spectral library-free (directDIA+) workflow and the Uniprot Burmese python (*Python bivittatus*) proteome as database (UP000695026, May 2024), using the default parameters. For global proteome Carbamidomethyl (C) was used as fixed modification, and Acetyl (Protein N-term), Oxidation (M), and Deamidation (NQ) as variable modifications. For PTM analysis, the PTM workflow was used, including Phospho (STY), GlyGly (K), and Acetyl (K) respectively as variable modification and setting the PTM probability cutoff to zero. Spectronaut intensity data matrices were imported to Perseus (1.6.10.45) and transformed into PTM site tables using the peptide collapse plugin.

### Proteomics Data Analysis

The statistical analysis was done using the R software (R version 3.5.0, RStudio version 1.0.143) and the Proteomics Toolset for Integrative Data Analysis Shiny app (ProTIGY, Broad Institute). Output data from Spectronaut 18 was filtered to remove protein groups with more than 20% missing values and protein groups with less than two peptides for global proteome analysis. For PTM analysis, sites with 3 valid values in at least one group and localization probability above 0.5 were kept. Intensities were normalized by median subtraction, and missing values were imputed by random sampling of the lower end of the normal distribution (width 0.3 and downshift 1.8). Two-sample moderated t-test pair-wise comparison of the normalized log_2_ intensities was used to determine the significantly regulated proteins and PTM sites among groups with an FDR cutoff of 5%. GO Biological Process over-enrichment analysis was performed using Enrichr^86^. PTM set enrichment analysis of the phospho-proteomics data was performed using ptmSEA, as previously described^87^.

### Protein Structural Modeling and Annotation

AlphaFold3^88^ was used for protein structural predictions with *Python bivittatus* MYH15 and TTN sequences. The Burmese python MYH15 converter domain sequence was ascertained by sequence homology with *Homo sapiens* MYH7. The highest confidence level AlphaFold3 structures were used for downstream annotation of post-translationally modified residues. ChimeraX^89^ (version 1.10) was used to annotate feeding-regulated residues in MYH15 and TTN.

### Statistical Analysis and Presentation

Statistical significance cut-offs for the RNA sequencing and proteomics datasets were established using adjusted p-values after Benjamini-Hochberg false discovery rate correction. Comparisons of two groups were made using an unpaired, two-tailed t-test. Comparisons of more than two groups with a single variable were performed by one-way ANOVA. Comparisons of more than two groups and two independent variables were performed by two-way ANOVA. If statistical significance was reached, Sidak’s or Tukey’s post-hoc test for multiple independent pairwise comparisons was used, as detailed in the figure legends. Unless otherwise noted in the figure legends, a p-value or adjusted p-value of < 0.05 was considered statistically significant. Data presentation was performed using GraphPad Prism (version 10) and Adobe Illustrator.

## DATA AVAILABILITY

The raw RNA sequencing data were deposited at the National Center for Biotechnology Information (NCBI) (BioProject ID: PRJNA1335253) and are publicly available. The raw proteomics data were deposited at ProteomeXchange using the PRIDE partner repository (Dataset ID: *will be made public upon acceptance*) and are publicly available. All other data associated with this study can be found in the text, figures, figure legends, or supplementary material. Requests for information or materials associated with this study should be directed to the corresponding authors.

## ACKNOWLEDGEMENTS

This study was supported by the Fondation Leducq Transatlantic Network of Excellence (21CVD02 to MG and LAL), the United States National Institutes of Health (F32HL170637 to TGM), and the American Heart Association (24PRE1195130 to DRH). We thank Dr. Jack Gugel, Skip Maas, Marcus Mullen, and Brooke Shepard for valuable scientific discussions and feedback on this study. We also acknowledge Aaron Rothchild and the staff in the Office of Animal Resources at CU Boulder for assistance with python colony husbandry. We thank the Shared Instruments Pool (RRID: SCR_018986), Department of Biochemistry, University of Colorado Boulder for the use of the Amersham Typhoon 5 imager (funded by NIH S10OD034218). We thank the BioFrontiers Computing Facility at the University of Colorado Boulder for High Performance Computing and data storage resources supported by BioFrontiers IT.

## AUTHOR CONTRIBUTIONS

Conceptualization: TGM, LSA, YT, MG, PM, and LAL. Methodology: TGM, LSA, KCW, EGM, YT, DRH, and BEK. Investigation: TGM, LSA, KCW, EGM, YT, DRH, BEK, LN, JL, and IL. Formal Analysis: TGM, LSA, KCW, EGM, DRH, and JL. Visualization: TGM, LSA, DRH, and JL. Funding Acquisition: TGM, DRH, JO, MG, and LAL. Resources: KSA, MG, PM, and LAL. Project Administration: MG, PM, and LAL. Supervision: KSA, JO, MG, PM, and LAL. Writing – Original Draft: TGM. Writing – Review and Editing: LSA, KCW, EGM, YT, DRH, BEK, JL, IL, LN, KSA, JO, MG, PM, and LAL.

## ADDITIONAL INFORMATION

This article includes Supplemental Data.

## CORRESPONDENCE

Requests for materials should be addressed to Michael Gotthardt, Philipp Mertins, or Leslie Leinwand.

