## Supplemental Information for "Multi-omics Insight into Cardiac Myofibril Remodeling in Post-Prandial Burmese Pythons"

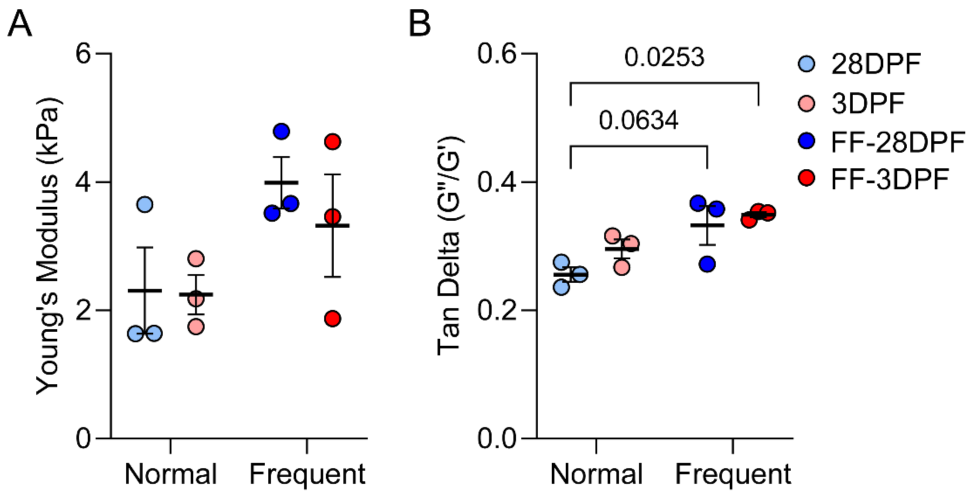


**Fig. S1 | Frequent feeding increases ventricular tissue stiffness and decreases elasticity. A.** Young's modulus of python ventricular tissue measured by oscillatory shear rheology; n = 3 biological replicates/group; two-way ANOVA with Tukey's post-hoc test; interaction: p = 0.617, row factor: p = 0.045, column factor: p = 0.545. **B.** Ratio of viscoelastic loss moduli (G'') to storage moduli (G'), representing the Tan delta, as determined by shear rheology; n = 3 biological replicates per group; two-way ANOVA with Tukey's post-hoc test; interaction: p = 0.532, row factor: p = 0.007, column factor: p =0.152.

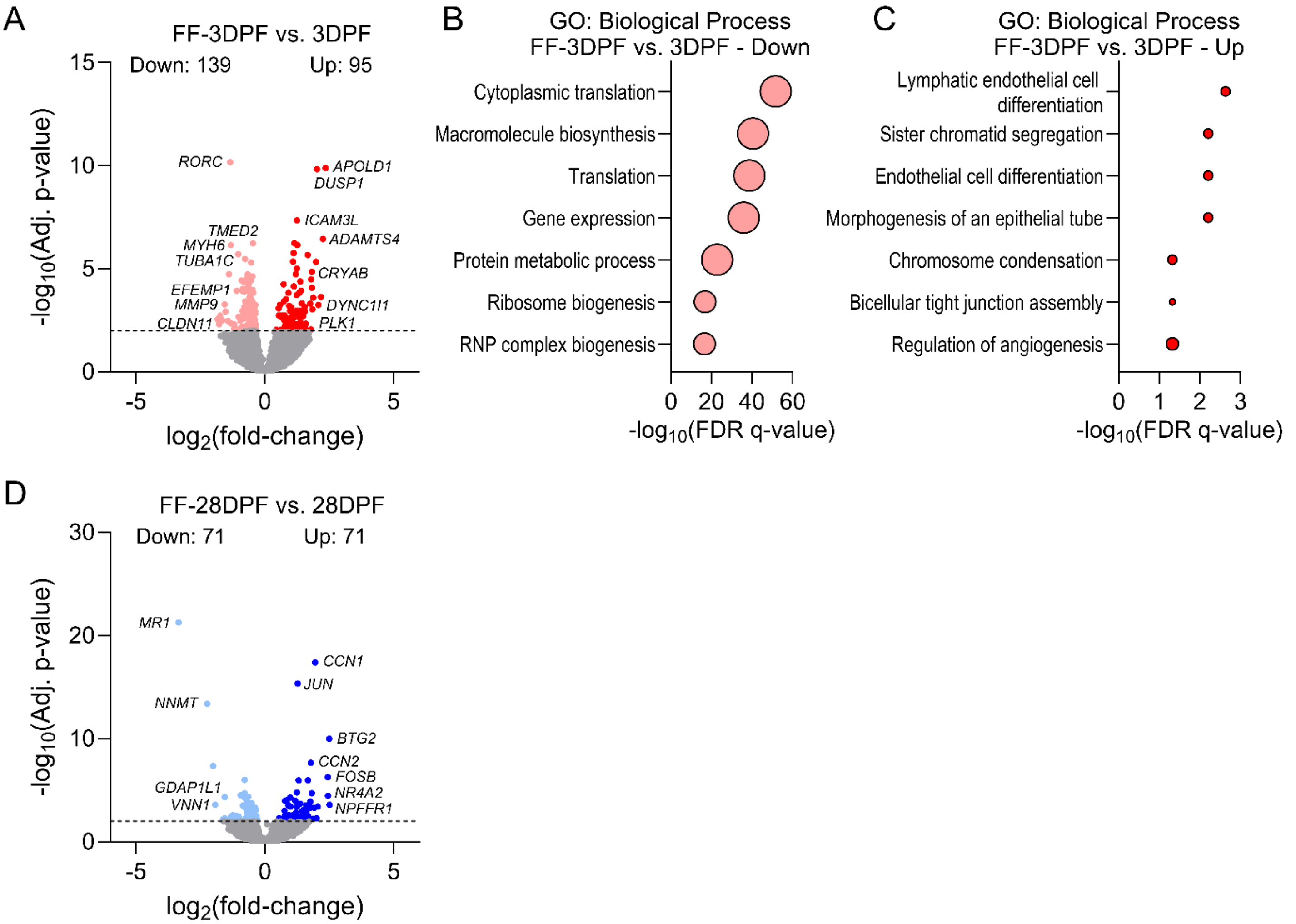


**Fig. S2 | Cardiac ventricular gene expression signatures associated with frequent feeding. A.** Volcano plot depicting fold-change, statistical significance, and number of significantly differentially expressed genes in the FF-3DPF vs. 3DPF comparison. **B-C.** GO: Biological Process enrichment for the downregulated (B) and upregulated (C) genes in the FF-3DPF vs. 3DPF comparison. **D.** Volcano plot depicting fold-change, statistical significance, and number of significantly differentially expressed genes in the FF-28DPF vs. 28DPF comparison. No GO: Biological Processes were identified as significantly enriched in this comparison.

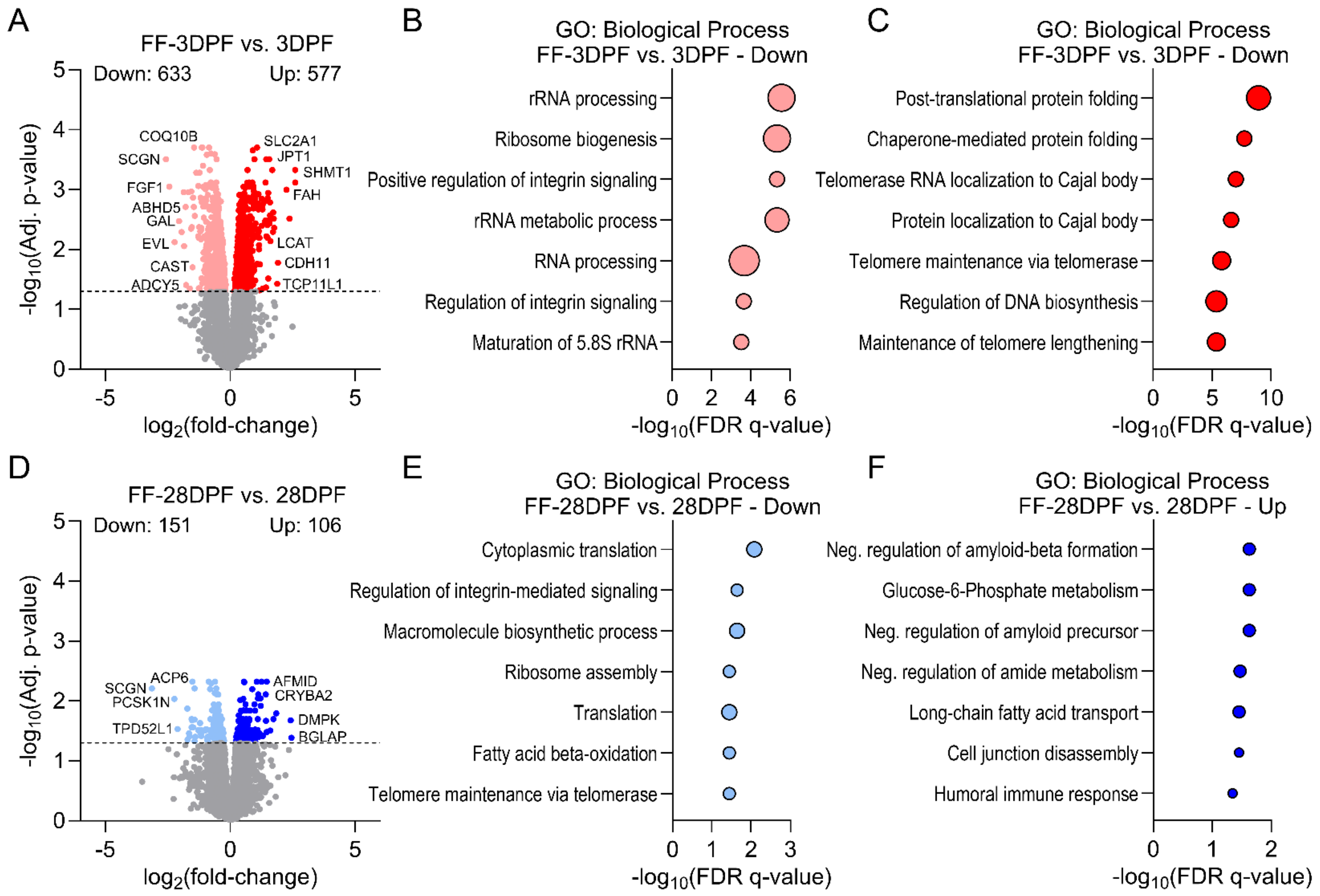


**Fig. S3 | Cardiac ventricular proteome signatures unique to frequent feeding. A.** Volcano plot depicting fold-change, statistical significance, and number of significantly differentially expressed proteins in the FF-3DPF vs. 3DPF comparison. **B-C.** GO: Biological Process enrichment for the downregulated (B) and upregulated (C) proteins in the FF-3DPF vs. 3DPF comparison. **D.** Volcano plot depicting fold-change, statistical significance, and number of significantly differentially expressed proteins in the FF-FF-28DPF vs. 28DPF comparison. **E-F.** GO: Biological Process enrichment for the downregulated (E) and upregulated (F) proteins in the FF-28DPF vs. 28DPF comparison.

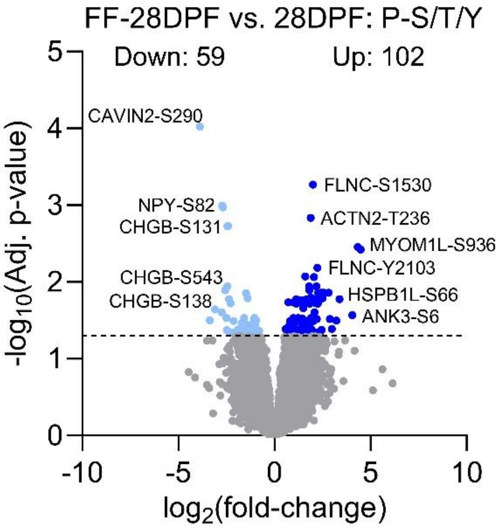


**Fig. S4 | Frequent feeding induces long-term phospho-proteome remodeling.** Volcano plot depicting fold-change, statistical significance, and number of significantly differentially expressed phospho-sites in the FF-28DPF vs. 28DPF comparison.

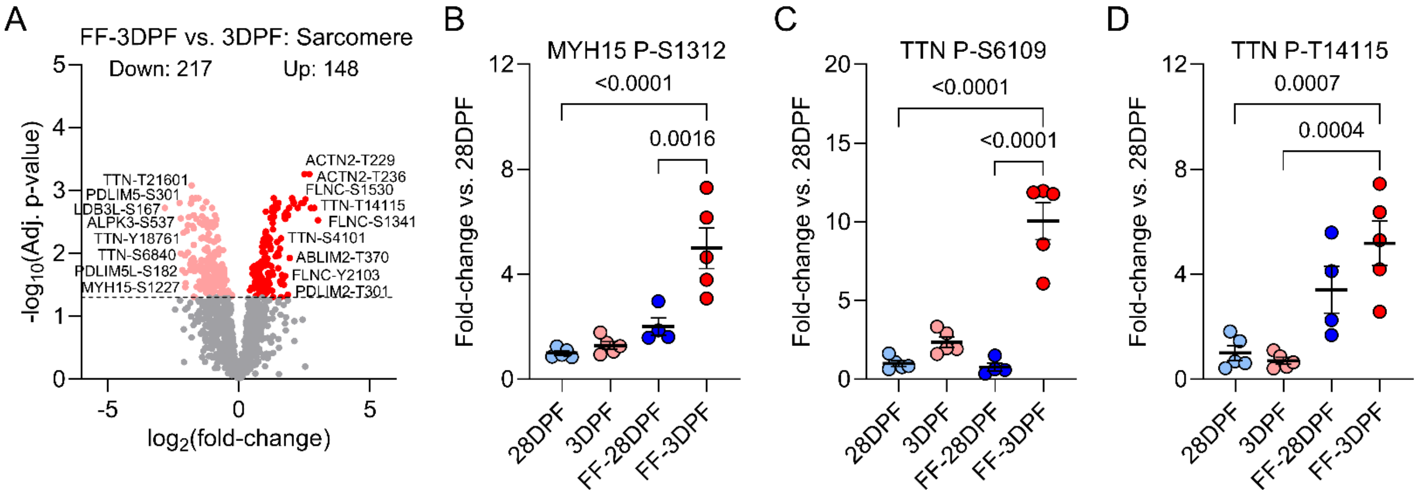


**Fig. S5 | Sarcomere protein phosphorylation changes induced by frequent feeding. A.** Volcano plot depicting fold-change, statistical significance, and number of significantly differentially expressed phospho-sites for sarcomere proteins in the FF-3DPF vs. 3DPF comparison. **B-D.** Notable phospho-sites that were significantly increased uniquely in frequent feeding; two-way ANOVA, Tukey post-hoc; B, interaction: $p = 0.008$, feeding frequency: $p < 0.0001$, time after feeding: $p = 0.002$; C, interaction: $p < 0.0001$, feeding frequency: $p < 0.0001$, time after feeding: $p < 0.0001$; D, interaction: $p = 0.108$, feeding frequency: $p < 0.0001$, time after feeding: 0.243; data are presented as the mean ± SEM.
